# Movement-related activity in the internal globus pallidus of the parkinsonian macaque

**DOI:** 10.1101/2024.08.29.610310

**Authors:** Daisuke Kase, Andrew J. Zimnik, Yan Han, Devin R. Harsch, Sarah Bacha, Karin M. Cox, Andreea C. Bostan, R. Mark Richardson, Robert S. Turner

## Abstract

Although the basal ganglia (BG) plays a central role in the motor symptoms of Parkinson’s disease, few studies have investigated the influence of parkinsonism on movement-related activity in the BG. Here, we studied the perimovement activity of neurons in globus pallidus internus (GPi) of non-human primates during performance of a choice reaction time reaching task before and after the induction of parkinsonism by administration of 1-methyl-4-phenyl-1,2,3,6-tetrahydropyridine (MPTP). Neuronal responses, including increases or decreases in firing rate, were equally common in the parkinsonian brain as seen prior to MPTP and the distribution of different response types was largely unchanged. The slowing of behavioral reaction times and movement durations following the induction of parkinsonism was accompanied by a prolongation of the time interval between neuronal response onset and movement initiation. Neuronal responses were also reduced in magnitude and prolonged in duration after the induction of parkinsonism. Importantly, those two effects were more pronounced among decrease-type responses, and they persisted after controlling for MPTP-induced changes in the between-trial variability in response timing. Following MPTP the trial-to-trial timing of neuronal responses also became uncoupled from the time of movement onset and more variable in general. Overall, the effects of MPTP on temporal features of GPi responses were related to the severity of parkinsonian motor impairments whereas changes in response magnitude and duration did not reflect symptom severity consistently. These findings point to a previously underappreciated potential role for abnormalities in the timing of GPi task-related activity in the generation of parkinsonian motor signs.

**New & Noteworthy:** Although the globus pallidus internus (GPi) plays a central role in the cardinal symptoms of Parkinson’s disease (PD), how parkinsonism alters the movement-related activity of GPi neurons remains understudied. Using a monkey model of PD, we found that: 1) the timing of GPi responses became uncoupled from movement onset. And 2) responses, especially decrease-type responses, became attenuated and prolonged. These abnormalities in GPi perimovement activity may contribute to the generation of parkinsonian motor signs.

## INTRODUCTION

What abnormalities in brain physiology underpin the motor impairments of Parkinson’s disease (PD)? One key step toward answering that question came with the discovery that lesions or stimulation of the posterior globus pallidus internus (GPi, primary output nucleus for the skeletomotor basal ganglia (1)) reduce the cardinal motor signs of PD, particularly akinesia, bradykinesia, and rigidity (2–5). That observation, coupled with the fact that dopamine replacement medication is also highly effective against those symptoms (6, 7), positions the GPi as a key anatomic link in the pathophysiologic cascade that leads from dopaminergic denervation to culminate in the manifestation of symptoms. Substantial uncertainty persists, however, regarding the specific abnormalities in GPi physiology that lead to parkinsonian symptoms.

Most pathophysiologic models of parkinsonism focus on the potential roles of abnormal tonic activity in GPi neurons in subjects at rest. For example, the classical “rate model” hypothesizes that an elevated tonic firing rate of GPi neurons leads to excessive inhibition of GPi-recipient motor centers (8, 9). An alternate group of models predict that abnormal patterns of GPi activity (e.g., bursty rhythmic activity and increased synchronization of firing between neurons) play a key role in pathophysiology (10–13). An unanswered question for these models is how the abnormalities in resting neuronal activity are translated into a disruption of phasic motor commands, and thereby, impairments in active movement. Abnormalities in motor command-related single-unit activity are indeed common in BG-recipient regions of the thalamus (14), midbrain (15–17) and motor cortex (18–20). The pathophysiologic models cited above assume that the translation from abnormal tonic discharge to disrupted motor commands occurs in BG-recipient thalamo-cortical, midbrain, or brainstem circuits, after the activity has been transmitted out of the BG (21–23) (23).

A seldom considered alternative is that abnormalities in the dynamic perimovement changes in GPi activity contribute directly to the disruption of motor commands in parkinsonism. Perimovement modulations in neuronal activity are common in the GPi of neurologically-normal animals (24–28). Given that GPi projections have a strong inhibitory influence on their targets (29), abnormal perimovement activity in GPi could have a direct disruptive influence on the generation of motor commands in GPi-recipient thalamocortical, midbrain, and brainstem motor circuits. Surprisingly, only one known study describes the activity of GPi neurons around the time of active movement in parkinsonian animals. Leblois et al. (30) reported that induction of parkinsonism in non-human primates (NHPs) led to an overall increase in the responsiveness of GPi neurons to active movement, a relative reduction in the incidence of perimovement decreases in firing (compared to that of increases), and shifts in the timing and duration of perimovement activity. Of particular note, perimovement activity shifted to *earlier* than normal onset times relative to movement onset and, in a subset of animals, earlier onset times relative to the sensory go-cue. That result does not align well with the common observation that go-cue –to-movement onset delays (i.e., reaction times, *RTs*) are increased in parkinsonian subjects (31–33) or with the general idea that parkinsonian bradykinesis reflects a general slowing of brain processes (34, 35). Leblois et al. also found that the duration of GPi perimovement responses was markedly prolonged after the induction of parkinsonism.

Some of those observations (30) are consistent with previously-proposed pathophysiologic mechanisms. For example, the increased responsiveness of GPi neurons to active movement fits with the general idea that parkinsonism is associated with a breakdown in the functional segregation between circuits throughout the BG (i.e., reduced neuronal selectivity for specific sensory stimuli) (36–39). Also, Leblois et al.’s observation that perimovement decreases in firing become less common in parkinsonism could be seen as consistent with Nambu et al.’s (40) dynamic version of the classical rate model, in which under-activation of the GPi-inhibiting direct pathway (the “pro-kinetic” pathway) and over-activation of GPi-exciting indirect pathways (“anti-kinetic” pathways) work together to yield deficient perimovement decreases in GPi activity and concommitent inadequate disinhibition of GPi-recipient motor circuits (41). No established model, however, predicts Leblois et al.’s finding of a shift in the latency of GPi perimovement activity to earlier than normal onset times.

Here, we sought to replicate and expand upon the results of Leblois et al. (30) with a specific aim to gain better understanding of how induction of parkinsonism changes the magnitude and timing of GPi perimovement activity. We used a choice reaction time (*RT*) reaching task, variations of which have been used in previous studies of parkinsonism in non-human primates (18, 20). During task performance, single-unit neuronal activity was sampled from the skeletomotor territory of the GPi before and after the induction of parkinsonism by intracarotid infusion of the dopamine-specific pro-toxin MPTP. The distribution of different types of perimovement changes in activity (i.e., increases vs. decreases) and detailed parameters (latency, magnitude and duration) of perimovement changes in activity were compared between pre– and post-MPTP states. We used a trial-by-trial analysis of response timing and duration to disambiguate measures that are typically confounded in standard mean across-trials analyses. Our trial-by-trial analysis also provided new insights into the effects of parkinsonism on the trial-to-trial variability of response timing and the temporal linkage of neural responses to different task events.

We found perimovement responses in a significant proportion of GPi neurons after the induction of parkinsonism. Trial-by-trial analysis of responses revealed a reduction in response magnitude and a prolongation of response duration in the de-jittered response. These effects of MPTP were exaggerated in decrease-type responses. Additionally, there was an increase in the variability of response onsets between trials. Notably, the temporal relationship between GPi responses and task events was altered significantly. Following MPTP, GPi responses shifted to be more closely linked to the timing of the Go cue as opposed to that of movement. These changes in the temporal aspects of neural responses correlated positively with the severity of parkinsonism between animals, suggesting a potential role for abnormal response timing in the generation of parkinsonian motor signs.

## MATERIALS AND METHODS

### Animals

The subjects were three adult macaque monkeys (*Macaca mulatta*: G, female 7.1kg, age 8-10 years; I, female 7.5kg; age 10-13 years; L, female 7.0kg, age 15 years). Animal L was included only for the histological analysis. Some data collected from these animals prior to MPTP administration were used for a previous project (42). All aspects of animal care were in accordance with the National Institute of Health Guide for the Care and Use of Laboratory Animals (43), the Public Health Service Policy on the Humane Care and Use of Laboratory Animals, and the American Physiological Society’s Guiding Principles in the Care and Use of Animals. All experimental protocols were evaluated and approved by the Institutional Animal Care and Use Committee.

### Surgery

General surgical procedures have been described previously (44, 45). The chamber implantation surgery was performed under sterile conditions with ketamine induction followed by Isoflurane anesthesia. Vital signs (i.e. pulse rate, blood pressure, respiration, end-tidal pCO2, and EKG) were monitored continuously to ensure proper anesthesia. A cylindrical titanium recording chamber was affixed to the skull at stereotaxic coordinates to allow access to the arm-related part of the GPi in the right hemisphere via a parasagittal approach. The chamber was oriented parallel to the sagittal plane at an angle of 32°. The chambers and head stabilization devices were fastened to the skull via bone screws and methyl methacrylate polymer. Prophylactic antibiotics and analgesics were administered post-surgically.

On protocols.io, we have posted two step-by-step protocols relevant to the surgical procedures (craniotomy https://dx.doi.org/10.17504/protocols.io.n92ldmz48l5b/v1 and implantation of head fixation and recording chamber hardware https://dx.doi.org/10.17504/protocols.io.5jyl8p9yrg2w/v1).

### Behavioral task

The animals performed a choice RT reaching task that has been described previously (45, 46). This task was running on LabView software (National Instruments, RRID: SCR_014325) and the source code of the task is available on Zenodo (DOI: 10.5281/zenodo.11397910). In brief, the animal was seated in a monkey chair facing a vertical black panel on which two target LEDs were positioned at shoulder height 7 cm to the left and right of midline. The animal began a trial by placing the left hand on a metal ‘home-position’ bar positioned at waist height to the left of the left hip. Upon completion of a random duration start-position hold period (SPHP, 2-10 sec, uniform distribution randomized trial-to-trial) one of the target LEDs was illuminated (“go-cue”, left or right LED selected pseudo-randomly). The animal was required to move its hand from the home-position to the illuminated target LED within a 1 sec response window relative to the onset of the go-cue, and hold at the target position for 0.5-1.0 sec (uniform distribution). A drop of pureed food reward was delivered at the completion of successful trials via a sipper tube and a computer-controlled peristaltic pump. A trial was marked as an error and reward was not delivered if the animal failed to complete the hold period, reached to the incorrect target, or if the 1 sec response window was exceeded. The animal was allowed to return its hand to the home-position and initiate the next trial with no time restrictions. The presence of the hand at the home-position and at targets was detected by infrared proximity sensors (Takex, GS20N).

Some timing parameters of the task were adjusted following MPTP administration to make it easier for the parkinsonian animals to perform the task successfully. The maximum response window was expanded to 3 sec relative to onset of the go-cue, and the hold period at the target position was shortened to 0.1 sec.

In addition, parkinsonian animals often failed to return the affected left hand to the start position at the end of a trial, after reward delivery. A milder form of this phenomenon has been described previously(46, 47). In the present study, if an animal failed to initiate a return-to-home movement within 4 sec of reward delivery then the experimenter intervened and provided manual assistance to return the animal’s hand to the home-position. Both animals readily learned to accept this manual assistance.

### Electrophysiological Recording

To ensure that single-unit recordings was restricted to the GPi, we first mapped each recording track to locate the borders of the GPi relative to GPe and internal capsule using glass-insulated tungsten electrodes (0.5-1.0 M Ω, Alpha Omega Co.). The extracellular spiking activity of neurons in the GPi was collected using multiple glass-insulated tungsten electrodes (0.5-1.0 MΩ, Alpha Omega Co.) or 16-channel linear probes (0.2-1.0 MΩ, Plexon Inc.). Signals were amplified (4x), band-pass filtered (2 Hz – 7.5 kHz), digitized at 24 kHz (16-bit resolution: Tucker Davis Technologies), and saved to disk as continuous signals. When stable single-unit isolations were available in one or more channels, neuronal and behavioral data were collected while the animal performed the behavioral task.

On protocols.io, we have posted protocols relevant to electrophysiological recording (https://dx.doi.org/10.17504/protocols.io.bp2l6xx91lqe/v1) and recording chamber maintenance (https://dx.doi.org/10.17504/protocols.io.5jyl8ppm9g2w/v1).

### Administration of MPTP

After completion of observations in the normal state, a hemiparkinsonian syndrome was induced by injection of MPTP into the right internal carotid artery (ICA; 0.5 mg/kg, (48)). This model of parkinsonism was chosen to ensure that animals could maintain themselves and remain healthy for the months-long period of post-intoxication recording. Use of this model also increased the likelihood that animals would continue performing the operant task following intoxication (20). The ICA MPTP administration procedure was performed under general anesthesia (1-3% Isoflurane), and prophylactic antibiotics and analgesics were administered postsurgically.

We have posted on protocols.io the step-by-step protocol used to administer MPTP: (https://dx.doi.org/10.17504/protocols.io.dm6gp3zw5vzp/v1).

Monkey G received one ICA infusion of 0.33 mg/kg MPTP hydrochloride (Sigma M0896-10MG). In monkey I, stable parkinsonian signs appeared after two ICA infusions of MPTP (0.33 mg/kg) followed by four intra-muscular injections (0.33∼0.55 mg/kg). The severity of parkinsonism was assessed by: (1) the degree of slowing and appearance of other impairments in the performance of the reaching task (see Results); (2) reduced overall mobility of the animal in its home cage accompanied by a dramatic increase in the tendency to turn toward the more depleted right hemisphere (clockwise)(48); (3) a marked reduction in the density of dopamine terminals in the dorsolateral putamen and substantia nigra compacta as measured by tyrosine hydroxylase (TH) staining in *post mortem* tissue. Dopamine replacement therapy was not administered at any time during the post-MPTP data collection period. Post-MPTP recording sessions started >30 days after the last MPTP administration. The timelines of MPTP administration, data collection, and the task performance (RT, MD, and freezing rate) are summarized in Supplemental Figure S1.

### Behavior in home cage

The behaviors displayed by the animals in their home cage were documented using a digital video camera during multiple recording sessions performed before and after the final ICA administration of MPTP (30 to 60 minutes of video per session). For monkey G, 4 and 9 video recording sessions were performed prior to and following MPTP administration, respectively. For monkey I, 5 and 5 video recording sessions were performed pre– and post-MPTP, respectively. The dates of video recordings are indicated in Supplemental Fig. S1. During each video recording session, the subject was housed alone in one upper cubicle of the home cage and access of husbandry staff to the room was restricted. The video data were transferred to a server after collection and used to evaluate the impacts of MPTP administration on spontaneous behavior in the home cage.

### Histology

After the last recording session, each monkey was given a lethal dose of sodium pentobarbital and perfused transcardially with saline followed by 10% formalin in phosphate buffer and then glycerol. The brains were blocked in place in the coronal plane, removed, cryoprotected with sucrose, and cut into 50 μm sections. Sections at 0.5 mm intervals were stained with cresyl violet. Reconstruction of microelectrode recording locations relative to nuclear boundaries was aided by a merging of information across cresyl violet sections, projection of microelectrode mapping information relative to structural MRI images (49), and the location of nuclear boundaries defined in parasagittal planes from an online atlas (BrainMaps: An Interactive Multiresolution Brain Atlas; http://brainmaps.org [retrieved on 2016])(50).

Selected coronal sections were processed for immunohistochemistry to visualize tyrosine hydroxylase (TH) for documentation of the loss of dopaminergic terminals from the striatum and cell bodies from the substantia nigra pars compacta (SNc). We have posted on protocols.io the step-by-step protocol used for the immunohistochemical staining for TH (protocol: https://dx.doi.org/10.17504/protocols.io.4r3l2q1dql1y/v1; antibody: Novus Cat# NB 300-109, RRID: AB_350437). For each animal, TH-positive neurons in the VTA and SNpc were counted manually (by DRH and DK and confirmed by ACB) on two 50 um thick sections spaced 1 mm apart through the midbrain. We compared the counts in our MPTP-treated animals (I and G) to those in an age-matched control (animal L, 16 yo, F rhesus macaque). Sections were imaged using a Hamamatsu NanoZoomer S60 Digital slide scanner (RRID:SCR_023762). Outlines and estimated borders between VTA and SNpc were drawn on scanned images using CorelDraw Graphics Suite (RRID:SCR_014235).

### Analysis

#### Home cage behavior

The video records of home cage behavior were divided into multiple files each containing a 5-minute segment of the video. Those segments were assigned randomized filenames. Two raters, blinded to the experimental conditions and file naming convention, viewed each video segment and counted occurrences of two spontaneous behaviors: (1) scratching or grooming of the body, counted separately when left and right limbs were used (hand or foot), and (2) full 360-degree rotations of the body, categorized separately as counterclockwise (leftward for the subject) and clockwise (rightward). Raw counts were converted into rates (events per minute). To evaluate MPTP-induced changes in the lateral bias of these behaviors, we subtracted the rates of right-sided behaviors from those of left-sided behaviors.

#### Behavioral data

All error trials and outliers in task performance were excluded from further analyses. Error trials included reaches to the incorrect target and failures to reach the target within the allowed interval after the go cue (1 sec and 3 sec for pre– and post-MPTP respectively). Outlier trials were detected according to RT and movement duration (*MD*). The RT was defined as the interval between the go-cue presentation and subsequent movement onset, and MD as the interval between movement onset and target capture. Trials were classified as outliers if either behavioral metric exceeded a threshold of 6 median absolute deviations from the mean.

#### Single-unit data

The stored continuous neurophysiologic signals were high pass filtered (Fpass: 300Hz, Matlab FIRPM, MathWorks, RRID: SCR_001622). For the bulk of these recordings, candidate action potentials were thresholded and sorted manually using Off-line Sorter (*OLS*, Plexon Inc., RRID: SCR_000012). Using that method, clusters of similar-shaped action potentials were identified across multi-way 2-D projections of waveform metrics. Clusters were accepted as representing the spiking of well-isolated single-units only if the cluster’s waveforms were of a consistent shape, could be separated reliably from other clusters as well as from background noise across a large fraction of the recording session, and violated a minimum refractory period criterion (1.5 ms) in only 0.5% of the resulting inter-spike intervals.

A subset of the neuronal data were sorted using a custom semi-automated spike-sorting algorithm (*TomSort*). The TomSort algorithm was used to identify single unit clusters in the neuronal recordings that were collected with 16-channel laminar probes in animal I following MPTP administration. After putative single unit clusters were identified by TomSort, those results were curated manually using OLS and the same acceptance criteria as applied when sorting was performed using OLS alone. A post-hoc comparison of results from the application of OLS and TomSort methods on the same recordings showed that TomSort identified nearly all of the single-units isolated using OLS alone. For example, across two recording sessions, TomSort successfully isolated 12 out of the 13 GPi units that were identified using OLS alone. But TomSort identified additional single-unit clusters that were not identified using OLS alone. Post-hoc inspection revealed that waveforms from those additional clusters were evident in the raw neuronal recordings, but those units were overlooked during manual sorting using OLS alone. The MATLAB code for TomSort is available on Github: (https://github.com/turner-lab-pitt/TomSort; DOI: 10.5281/zenodo.11176979).

We submitted all isolated units to a procedure that flagged instances of anomalous spike waveform features. Such instances may indicate, for example, that a cluster of waveforms did not originate in the soma of a GPi cell. Briefly, we summarized the median unit waveforms according to three features (51–54): the half-width, the difference in the peak and trough magnitudes (normalized by waveform amplitude), and the difference in the pre– and after-hyperpolarization peaks (normalized by their sum). For each MPTP state (and collapsed over the two subjects), we represented each unit as a point in a three-dimensional feature space, which was in turn submitted to the MATLAB implementation of DBSCAN (55). DBSCAN identifies outliers as points that lack neighbors that fall within a pre-defined distance. We discarded all outlier units, resulting in the exclusion of 9.40% and 3.17% of the units isolated pre– and post-MPTP, respectively. The MATLAB code for this analysis is available on Github (https://github.com/turner-lab-pitt/outlier-waveform-detection; 10.5281/zenodo.11118235), and input data for this code are available on Zenodo (DOI: 10.5281/zenodo.15276457).

After removing the units with outlier spike waveform features, we defined acceptable ranges of firing rates for pre– and post-MPTP populations separately. We calculated mean firing rates across whole recording periods for each unit population from both animals and then obtained the median and median absolute deviation (MAD) across those populations. The range of acceptable firing rates was defined as rates that fell within 1.5 × MAD of the median firing rate (22.9−117.7 Hz for pre-MPTP units and 1.5 −76.0 Hz for post-MPTP units). Single units were accepted for further analysis if their activity met the spike sorting, waveform shape and firing rate criteria described above and their activity was sampled over at least 10 trials for each target direction in the session.

#### Discharge rate and pattern at rest

The effects of MPTP intoxication on resting neuronal activity were quantified using standard methods (56, 57). Those analyses were applied to spike trains during the start-position hold periods (SPHPs) of the behavioral task – intervals of unpredictable duration (2-10 sec) during which an animal held its left hand stationary at the start position while awaiting onset of the go-cue. A neuron’s mean firing rate was calculated as the total number of spikes detected across all hold periods divided by the summed duration of all hold periods. Episodes of burst firing – discrete period of markedly elevated firing rate – were detected using the Poisson Surprise Method (57, 58). Bursts were defined as groups of 4 or more spikes whose inter-spike intervals (ISIs) were unusually short compared with other ISIs of the same spike-train. We used a surprise threshold of 5, which equates to p<0.05 that the candidate burst would occur as a part of a Poisson-distributed sequence of spikes. The overall “burstiness” of a cell was quantified as the fraction of spikes that occurred during bursts relative to the total number of spikes found across all SPHPs.

Rhythmic modulations in firing rate were detected using a “shuffled normalization” method (59–61). The discrete Fourier transform (FFT) was applied to non-overlapping 512-ms long segments of a spike train’s delta function smoothed with a Hanning window of the same length. (These spike train segments were extracted separately from each SPHP to avoid possible anomalies due to data discontinuities at SPHP concatenation boundaries. SPHPs containing fewer than four spikes were excluded from this analysis.) The resulting “primary” spectral density estimate (0–500-Hz, 2-Hz resolution) was normalized by dividing it by a “control” spectrum. The control spectrum was the mean of 100 spectra computed after 100 shufflings of the same ISIs as used to compute the primary spectrum. This normalization compensated for distortions in spectral estimates attributable to a neuron’s refractory period and thereby improved the detection of low frequency oscillations (60). The shuffle-normalization procedure yielded spectra that varied around a normalized value of 1. Peaks in the normalized spectrum between 4-Hz and 100-Hz were tested for significance relative to the SD of the spectrum in the 150–250-Hz “control” range. The omnibus threshold for significance (p=0.05) included a Bonferroni correction for multiple comparisons (0.05 / 51 spectral points tested between 4-Hz and 100-Hz; actual threshold p<9.8×10^-4^). If a spectral peak exceeded the threshold at more than one frequency bin, then the central rhythmic frequency was defined as the spectral bin with the highest power.

#### Perimovement discharge

We tested for perimovement changes in single-unit spike rate using a method described previously (45). First, spike density functions (SDFs) were constructed by convolving each unit’s spike time stamps (1 kHz resolution) with a Gaussian kernel (σ = 25 ms). Spike trains were aligned to the time of movement onset in each trial, and across-trial mean SDFs were constructed separately for reaches to each target. Significant modulations in the mean SDF were tested for during a test window that extended from the across-trials median time of go-cue onset to the across-trials median time of movement offset. The significance of modulations during that test window was determined relative to baseline defined as the mean and standard deviation of the mean SDF across a 700 ms window that ended at the beginning of the test window (defined above), after correcting for linear trends in the mean SDF over the baseline window. We defined a peri-event response as a statistically significant elevation or depression in the mean SDF that lasted at least 60 ms relative to baseline (e.g., Fig. 3A, solid vertical lines; t-test, one-sample versus baseline; omnibus p < 0.001 after Bonferroni correction for multiple comparisons). Significant perimovement modulations in mean firing rate were classified as increase-or decrease-type “responses.” As described previously (62), the perimovement modulation of some single-units was polyphasic, being composed of one or more significant increase and decrease in firing rate during the test window. Those responses were classified according to the sign of the earliest significant modulation – either polyphasic increase/decrease or polyphasic decrease/increase [referred to hereafter as “poly (+/−)” and “poly (−/+),” respectively].

#### Quantification of response metrics

The onset latency of a response was defined as the time of the earliest significant point in the response (1 ms resolution) relative to the alignment event of the mean SDF (Fig. 4A *vertical green line*). For monophasic increase– and decrease-type responses, the magnitude of a response was defined as the firing rate maximum or minimum, respectively, relative to the unit’s baseline firing rate estimated at the detected time of response onset (Fig. 4A). The duration of a response was defined as the full-width of the response at half of its maximum change in firing rate [i.e., full-width at half-max (*FWHM*), or at half-minimum for decrease-type responses]. For polyphasic responses, response magnitude and duration metrics were computed as outlined above, but only for the initial phase of the response.

We also tested for effects of movement direction (i.e., between movements to left and right targets) on a neuron’s perimovement firing rate. Two approaches were used. First, following an established method (42), trial-by-trial spike counts from a 300 ms window starting at a unit’s earliest detected response onset were compared between left– and right-target trials. If a unit’s spike rate differed significantly between the two directions (p<0.05; Matlab RANKSUM) then that unit’s perimovement activity was considered to be “directional.” A second approach was used to control for potentially confounding effects of alterations in task performance with the induction of parkinsonism. The second approach took trial-by-trial spike counts from the whole response interval of each trial (starting at go-cue onset and ending at movement termination) and compared them between left– and right-target trials. A unit’s response period activity was considered directional if it differed significantly between the two movement directions (p<0.05; Matlab RANKSUM).

#### Trial-by-trial detection of response onsets

Measures of response magnitude and duration taken from a mean across-trials SDF are susceptible to distortions that depend on how variable the timing of the response is from trial to trial (i.e., its “temporal jitter”). For example, in a comparison of across-trials mean SDFs, what appears to be reduction in response magnitude and increase in duration could in fact be attributable to a simple increase in the trial-to-trial variability of response timing (schematized in Fig. 4F-G). In other words, the net effect of increased trial-to-trial jitter in response timing is indistinguishable from a true reduction in response magnitude and increase in response duration when measurements are taken from a mean across-trials SDF.

To resolve this ambiguity, we detected the onsets of neuronal responses on a trial-by-trial basis using a general approach that has been described in detail elsewhere (62, 63). First, only units found to have a significant perimovement response in the mean across-trials SDF were subjected to this analysis. Among those units, the type of response detected in the mean SDF was used to guide the subsequent trial-by-trial analysis. (The approach used for monophasic increases is detailed here.) Second, perimovement SDFs for individual trials were computed by convolving spike time stamps (1 ms resolution) with a Gaussian kernel (σ = 25 ms). Third, for each single trial SDF, we prepared a 200-point (200 ms) response kernel consisting of a simple step function in which the initial 100 points were set to the minimum firing rate in the single-trial SDF and the second 100 points are set to the maximum firing rate. Fourth, the quality of fit between the kernel and segments of the single-trial SDF were computed while shifting the kernel at millisecond steps from the time of go-cue onset to the end of movement. The time shift that resulted in the best fit (minimum summed squared error) between kernel and segment of single-trial SDF was taken to be the time of response onset for that single trial. This procedure was repeated for the single-trial SDFs from each valid behavioral trial. For decrease-type responses, we first inverted the single-trial SDFs and then processed as described above for increase-type responses.

We then used those trial-by-trial measures of response timing to: 1) analyze the temporal linkage of responses to different task events (e.g., neural activity time-locked to movement onset; approach detailed in *Results*), 2) estimate the influence of parkinsonism on the fundamental trial-to-trial variability of response timing, and 3) test for effects of parkinsonism on response metrics (magnitude and duration) after controlling for potentially confounding influences of differences in a response’s trial-to-trial temporal jitter.

For item 2) above, the fundamental variability of response timing was quantified as the residual trial-to-trial variability in response timing after controlling for potential influences of RT variability and the temporal linkage of the response to specific task events. First, the linear relationship between trial-by-trial response latency and RTs was determined by linear regression (Matlab, FITLM) to model the RT-dependent variability in response timing. Residuals from that regression were taken to reflect the fundamental trial-to-trial variability in response timing. The interquartile range (IQR) of those residuals was used as a measure of the trial-to-trial dispersion in response timing.

We also used the trial-by-trial estimates of response timing to estimate “true” response metrics (magnitude and duration) after controlling for the potentially confounding influences of differences in response temporal jitter. For each response, we removed the effects of temporal jitter by realigning single-trial SDFs to the response itself. First, in individual single-trial SDFs, we found the earliest maximum change in firing rate (Matlab, FINDPEAKS; firing rate maximum for increase-type responses and minimum for decrease-type responses) that occurred following the previously determined time of response onset for that trial. Second, whole single-trial SDFs were shifted in time so that the times of peak change aligned across trials. Third, those realigned single-trial SDFs were averaged across-trials to yield a response-aligned mean SDF. Finally, response metrics (magnitude and duration) were extracted from the resulting response-aligned mean SDF using methods described above under *Quantification of response metrics*.

#### Statistics

##### ANOVA

We tested for potentially interacting effects of parkinsonism, animal, direction of movement and response type (e.g., increase-versus decrease-type responses) using two– and three-way analyses of variance (MatLab, ANOVAN). Tukey’s test was used for post-hoc analyses of the ANOVA results.

##### Standardized residuals analysis

When results from a chi-square test rejected the null hypothesis, we isolated the source of the statistically significant effect using a standardized residuals approach (64, 65).

##### Interquartile range

The dispersion of response onset times was quantified using the interquartile range (*IQR*), which is defined as the range that encompasses the central 50 percent of the data (66).

##### Clustering analysis

Distributions were tested for uni-versus poly-modal shapes using the Bayesian information criterion (*BIC*). BICs were obtained by fitting Gaussian mixture models to a distribution, considering a range of cluster numbers from one to four in this study (MatLab, FITGMDIST). The optimal number of clusters was determined by selecting the number of clusters with the smallest BIC (67).

## RESULTS

### Effects of MPTP administration: behavior and histology

The animals were rendered moderately hemiparkinsonian following the administration of MPTP as evidenced by changes in home cage behavior that were consistent with parkinsonism. Overall mobility was reduced, movement was slowed, and the animals adopted a hunched posture with increased flexion of the left limbs (contralateral to the depleted hemisphere). The animals exhibited a marked reduction in their preference for using the limbs on the left side of their body. For example, blinded scoring of in-cage videos showed that the limb used to perform grooming movements shifted strongly toward the right side of the body following MPTP (Wilcoxon rank test; *p* < 0.01 for monkey G and *p* < 0.05 for monkey I; Supplemental Fig. S3A). Additionally, both animals showed an increased tendency to turn in the clockwise direction (toward the more-depleted right hemisphere). The rate of clockwise rotations, relative to counterclockwise rotations, increased in both animals (Wilcoxon rank test; *p* < 0.01 for both monkeys; Supplemental Fig. S3B). Rigidity was increased at joints of the left arm and leg. We found no evidence of tremor in either animal, consistent with previous observations that MPTP-treated macaques seldom exhibit frank parkinsonian tremor (48). Examinations performed at regular intervals throughout the post-MPTP data collection period supported the persistence of these signs (Supplemental Fig. S1).

Despite the presence of parkinsonian impairments, both animals were able to perform the choice RT reaching task, albeit at reduced levels of performance. Both animals showed significant slowing of RTs and of the durations of outward reaching movements (MDs) to capture the target (Fig. 1F-H; 3-way ANOVA main effect of MPTP; F(_1,35652_)=31802, p < 0.01 for RT and F(_1,34986_)=50749, p < 0.01 for MD). Although significant slowing was observed in both animals, the degree of slowing was much more severe in monkey G than in monkey I (3-way ANOVA effect of MPTP × animal interaction; F(_1,35649_)=39.0, p<0.01 for RT and F(_1,34983_)=291.5, p<0.01 for MD).

**Figure 1.**
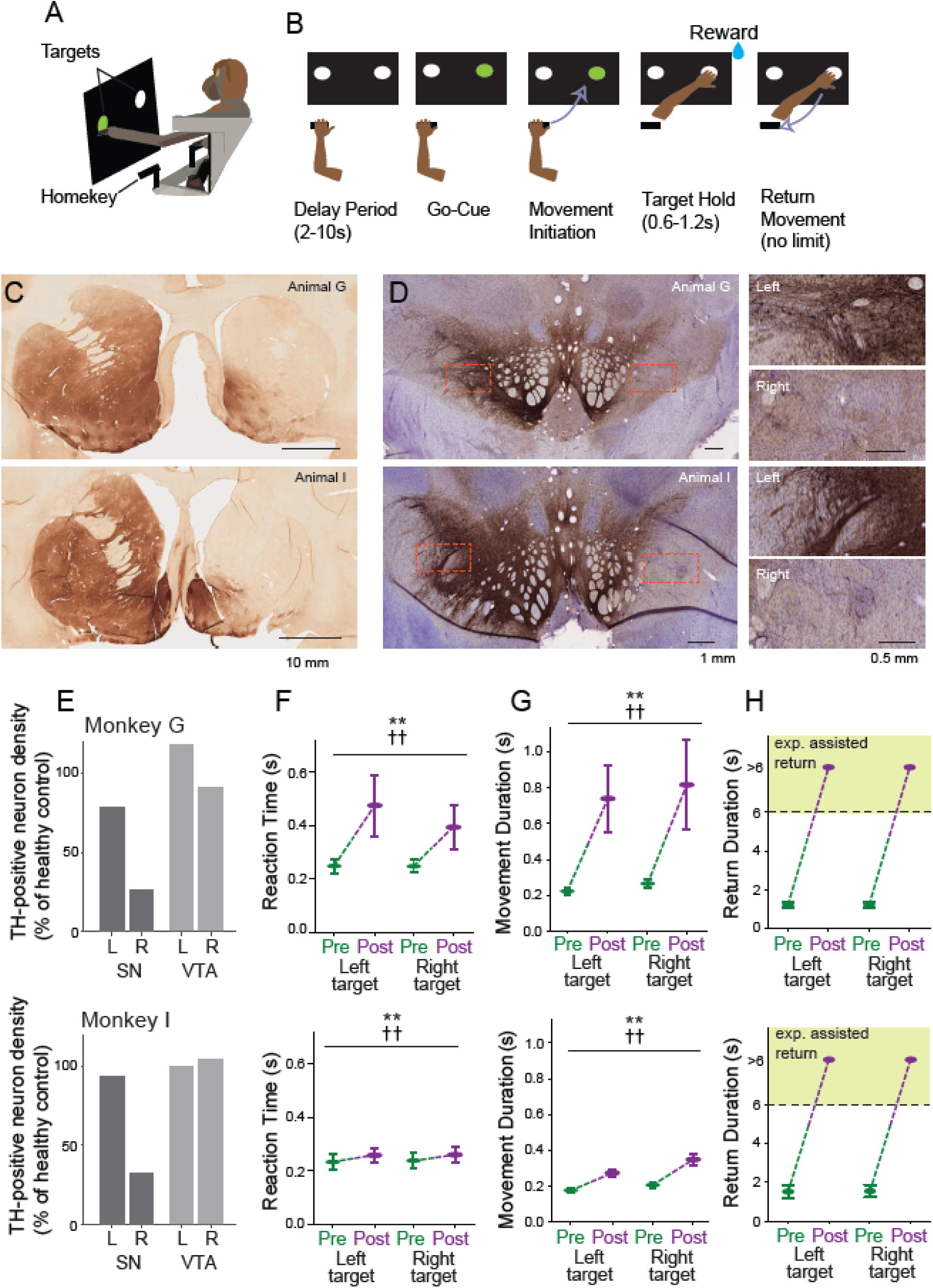
*A-B*: Schematic of task apparatus and the choice RT reaching task. *C*: Histologic immunolabeling for tyrosine hydroxylase (TH) revealed the depletion of dopaminergic innervation from the right striatum of both animal G (*top*) and I (*bottom*). *D*: Immunolabeling for TH (*brown*) and Nissl staining (*magenta*) demonstrate a marked reduction in TH labeling in the right lateral SNc of both animal G (*top left*) and I (*bottom left*). Note that TH labeling in the more medial right ventral tegmental area (VTA) was relatively preserved following the ICA MPTP administration. Inset panels (*right*) provide higher-magnification views of matching regions of the left and right lateral SNc (*red dashed rectangles*) and evidence the depletion of TH-label from the right lateral SNc. *E*: Densities of TH-positive somata in the SN (*black bars*) and VTA (*gray bars*), shown ad a percentage relative to the corresponding regions in the healthy control monkey L. Both animal G (*top*) and I (*bottom*) showed a selective depletion of TH-positive neurons in the right SN, while the left SN and both VTAs remained relatively preserved. *F-H*: Behavioral measures of task performance for both animals before and after MPTP administration (*green* and *magenta*, respectively). Both animals performed the task in a highly stereotyped fashion with short RTs (*E*), MDs (*F*), and return durations (*G*) prior to MPTP administration. RT and MD were slowed significantly following MPTP. *G*: Following MPTP, both animals displayed an extremely prolongation of the time required to return the hand to the start position. The experimenter intervened and assisted return movements if they were not initiated by the animal within 4 sec. of reward delivery, thus resulting in return durations of >6 sec (*green shading*). Means +/−SEM. Statistical differences were computed using a 3-way ANOVA followed by Tukey’s test; (**: main effect of MPTP, p<0.01; ††: MPTP × target interaction, p<0.01).

The MPTP-induced slowing of RTs and MDs was somewhat more severe for movements to the right target (located further away from the start position and ipsilateral to the lesioned hemisphere) as compared with movements to the left target (3-way ANOVA effect of MPTP × target; F(_1,35649_)=2.7, p<0.01 for RT and F(_1,34983_)=3.1, p<0.01 for MD). That target-dependent slowing was more severe in monkey G than in monkey I for RTs (3-way ANOVA effect of MPTP × target × monkey; F(_1,35649_)=2.3, p<0.01) but not for MDs (F(_1,34983_)=0.01, p=0.53).

Finally, during performance of the behavioral task, both animals displayed a severe impairment in the ability to return the hand to the start position after reward delivery. In the neurologically normal state, animals returned their left hand to the start position with minimal delay after reward delivery. Following MPTP, however, the animals rarely if ever initiated a return movement. Instead, the animal’s left arm remained extended with the hand held stationary near the target for seconds following reward delivery. Similar freezing-like arrests have been described previously in primate models of parkinsonism (46, 68). In order to collect adequate numbers of trials during post-MPTP recording sessions, if an animal did not return its hand to the start position within 4 seconds following reward delivery, the experimenter intervened and manually moved the animal’s left arm to bring the hand into contact with the start position. Thus, in our post-collection analysis of behavior, any return movements that lasted longer than 6 sec were classified as “experimenter assisted return” (Fig. 1H) and were considered to be categorically different from the rapid animal-initiated return movements of the pre-MPTP state. Experimenter-imposed manual assistance was required for essentially all of the trials collected following MPTP administration (chi-square test; *χ^2^*(1, n=35309) = 35309, p=0.000; see *Freezing Rate* in Supplemental Fig. S1).

In both animals, post mortem immunohistochemistry demonstrated a near complete loss of TH-positive neuronal elements from the right dorsal striatum (Fig. 1C) and a marked reduction in the density of TH-positive somata in the ventral lateral substantia nigra pars compacta (Fig. 1D). The density of TH-positive somata in the right substantia nigra (SN) was reduced to 26% of normal in monkey G and 32% of normal in monkey I, relative to densities found in a healthy control monkey (Fig. 1E). In contrast, TH-positive somata in the left SN and ventral tegmental area on both sides were relatively preserved. Representative images of immunohistochemical staining from control monkey L are shown in Supplemental Fig. S2 along with maps of the density of identified TH-positive neurons and tabular results of TH-positive somata counts.

### Neuronal database and activity at rest

A total of 371 single-units met the criteria to be included as GPi neurons (180 and 191 pre– and post-MPTP respectively). For recording sessions before MPTP administration, units were studied over the course of 136 ± 92 trials of the behavioral task (mean ± SD; mean 68 trials for each of two movement directions). Following MPTP administration, units were studied over the course of 100 ± 42 trials of the behavioral task (mean ± SD; mean 49 trials for each of two movement directions). Units were sampled from relatively wide-spread regions of the posterior GPi during both pre– and post-MPTP sampling periods (Fig. 2).

**Figure 2.**
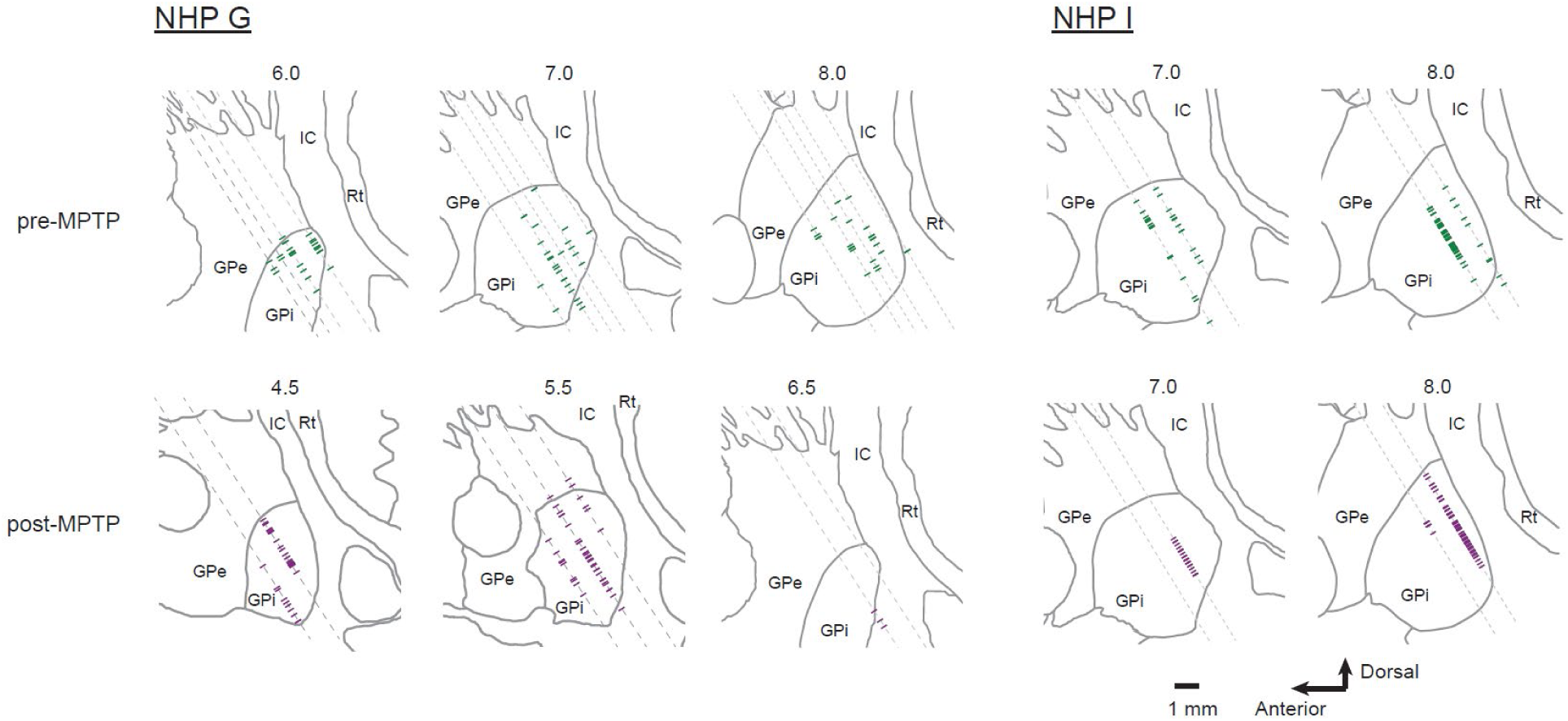
Reconstructed locations of all GPi single-units included in the database. Locations are projected onto parasagittal sections at 1 mm intervals separately for animal G and I, pre-(*green*) and post-MPTP (*magenta*). The line drawing delineating the GPi boundary was taken from a standard atlas (50), which was subsequently adjusted to align with the structural MRIs and microelectrode mapping results from individual animals.

The baseline activity of GPi neurons differed between pre– and post-MPTP periods in several ways, as would be expected with the induction of parkinsonism (10). Single-unit baseline firing properties were measured during a recording session’s start-position hold periods (SPHPs) – random duration (2-10 sec uniform distribution) periods of quiet attentive rest during which the animal held the left hand stationary at the start position in anticipation of go-cue presentation. Mean firing rates were reduced significantly post-MPTP (means ± SD: 69.0 ± 24.7 and 26.5 ± 19.2 Hz for pre– and post-MPTP respectively; Wilcoxon rank test; p<0.001, Table 1). Reductions in mean resting firing rates were found in both animals, but the magnitude of the reduction was more dramatic in monkey I (−50.69 sp/s) than in monkey G (−31.4 sp/s; 2-way ANOVA, MPTP × animal interaction; F(_1,367_)=19.7, p<0.01). The reduction in mean firing rate runs contrary to predictions of the classical model of PD pathophysiology (8, 9). The variability in inter-spike intervals was also elevated significantly post-MPTP (Wilcoxon rank test; p<0.001, Table 1).

**Table 1:** Activity at rest.

|  | NHP G |  | NHP I |  | Total |  |
| --- | --- | --- | --- | --- | --- | --- |
|  | pre-MPTP | post-MPTP | pre-MPTP | post-MPTP | pre-MPTP | post-MPTP |
| Number of Units | 92 | 82 | 88 | 109 | 180 | 191 |
| Firing rate (mean ± SD) | <u>68.6 ± 22.8</u> | <u>37.2 ± 21.2</u> | <u>69.3 ± 26.6</u> | <u>18.4 ± 12.7</u> | <u>69.0 ± 24.7</u> | <u>26.5 ± 19.2</u> |
| Coefficient of variation of ISI (mean ± SD) | <u>1.1 ± 0.4</u> | <u>1.3 ± 0.5</u> | <u>1.1 ± 0.5</u> | <u>1.4 ± 0.7</u> | <u>1.1 ± 0.5</u> | <u>1.4 ± 0.6</u> |
| Proportion of spikes in burst (mean ± SD) | <u>0.10 ± 0.11</u> | <u>0.20 ± 0.17</u> | <u>0.11 ± 0.12</u> | <u>0.20 ± 0.20</u> | <u>0.10 ± 0.12</u> | <u>0.20 ± 0.19</u> |
| Normalized Intra-burst firing rate (mean ± SD) | 4.7 ± 1.3 | 6.0 ± 6.4 | <u>4.4 ± 1.5</u> | <u>6.8 ± 9.7</u> | <u>4.6 ± 1.4</u> | <u>6.5 ± 8.4</u> |
| Oscillation at rest (units) | 47 (51%) | 33 (40%) | 28 (32%) | 23 (21%) | 75 (42%) | 56 (29%) |
| Theta (4-7.9 Hz) | **0** | **3 (8%)** | 0 | 1 (4%) | **0** | **4 (6%)** |
| Alpha (8-12 Hz) | **1 (2%)** | **4 (11%)** | 1 (3%) | 0 | 2 (2%) | 4 (6%) |
| Beta (12.5-30 Hz) | 27 (49%) | 24 (65%) | **5 (16%)** | **17 (65%)** | **32 (37%)** | **41 (65%)** |
| Low Beta (12.5-20 Hz) | 0 | 23 (92%) | 3 (60%) | 5 (29%) | 3 (9%) | 28 (67%) |
| High Beta (20.5-30 Hz) | 27 (100%) | 2 (8%) | 2 (40%) | 12 (71%) | 29 (91%) | 14 (33%) |
| Gamma (30.5-100 Hz) | **27 (49%)** | **6 (16%)** | **25 (81%)** | **8 (31%)** | **52 (60%)** | **14 (22%)** |
| Ramping of firing rate at rest (units) | 79 (86%) | 74 (90%) | 81 (92%) | 106 (97%) | 160 (89%) | 180 (94%) |
| Positive slope (units) | 32 (41%) | 33 (45%) | 35 (43%) | 50 (47%) | 67 (42%) | 83 (46%) |
| Negative slope (units) | 47 (59%) | 41 (55%) | 46 (57%) | 56 (53%) | 93 (58%) | 97 (54%) |

Underlined numbers indicate significant difference between pre- and post-MPTP (Wilcoxon rank sum test; single line $p < 0.05$ , double line $p < 0.01$ ). Bold numbers indicate significant difference between pre- and post-MPTP (Residual analysis following chi-square test, $p < 0.05$ )

Previous studies have also reported an increased tendency for GPi units to emit spikes within bursts (discrete periods of elevated firing rate) in parkinsonian animals (69, 70). In our dataset, the proportion of spikes that appeared within bursts increased markedly following MPTP administration (Wilcoxon rank test; p<0.05, Table 1). Both animals showed similar results when considered individually. Mean firing rates within bursts were reduced post-MPTP (means 296 and 122 spikes/sec pre– and post-MPTP, respectively; Wilcoxon rank test; p<0.01, Table 1). That result, however, was likely biased by the overall reduction in mean baseline firing rates post-MPTP (described above). When we normalized each unit’s intra-burst firing rates relative to its baseline firing rate, the mean normalized values increased significantly between pre– and post-MPTP periods (Wilcoxon rank test; p<0.05, Table 1).

Another often-observed neural correlate of parkinsonism in primates (human and non-human) is an increase in the prevalence of rhythmic modulations in firing rate (“oscillatory activity”) across a range of low frequencies (8-30 Hz; alpha and beta frequency bands) (71, 72). We used a shuffled normalization spectral analysis (60) to detect oscillatory spiking activity during SPHPs. The incidence of spectral peaks in the beta frequency range increased significantly following the induction of parkinsonism (standardized residual analysis; p<0.001, Table 1). Similar increases were found in the data from each animal considered individually, although that change reached significance only in NHP I (standardized residual analysis; p<0.001 and p=0.11 for NHPs I and G respectively). The increase in beta band oscillatory activity was accompanied by a significant reduction in the prevalence of spectral peaks in the gamma frequency range (standardized residual analysis; p<0.001). Since baseline neuronal firing rates were reduced post-MPTP placing them closer to the beta band range than firing rates pre-MPTP, it is possible that the increased prevalence of beta band oscillatory activity post-MPTP was an indirect product of the reduction in firing rates. To rule out that potential bias, we repeated the analysis using a subgroup of GPi units that had firing rates between 30 and 50 Hz during SPHPs. Within this subgroup, mean firing rates did not differ between pre– and post-MPTP populations (mean ± SD: 40.6 ± 6.4 Hz pre-MPTP vs. 38.6 ± 6.3 Hz post-MPTP; Wilcoxon rank test, p = 0.06). However, beta band oscillatory activity was more common in the GPi units collected post-MPTP (18.1% pre-MPTP vs. 34.7% post-MPTP; chi-square test, χ²(1, n=211) = 7.6, p < 0.01). Thus, beta band oscillatory activity was more common post-MPTP and that was not solely attributable to the reduction in firing rates during the same period.

To summarize, several aspects of GPi baseline activity changed following the administration of MPTP. The observed increases in firing variability, burst firing and beta frequency oscillatory activity were all consistent with a large literature describing neurophysiologic hallmarks of parkinsonism (10, 73). Although the observed reduction in mean firing rates runs contrary to the classical model of parkinson’s pathophysiology (8, 9), two factors should be kept in mind. First, nearly all past studies examined the effects of parkinsonism on GPi activity in animals that were not engaged in a defined behavioral task. Here, we studied GPi activity during the SPHPs of a well-learned task. GPi activity during those two distinct behavioral states may be affected differently by the induction of parkinsonism. Second, numerous studies have reported that the mean firing rate of GPi neurons (or SNr neurons in the rodent) is not a reliable predictor of the severity of parkinsonian symptoms (69, 73, 74). Overall, the changes in GPi baseline activity described here following MPTP administration were consistent with the animals’ parkinsonian status.

Unit activity during SPHPs also frequently showed small but significant ramp-like changes in firing rate (Fig. 3A; linear regression; p<0.05). This observation, described previously for neurologically normal animals (42), is likely related to the progressively increasing probability (i.e., the hazard rate) of go-cue onset when, as in our task, SPHP durations are distributed uniformly (75, 76). Ramping activities were found in large fractions of the GPi units sampled both before and following MPTP administration (89% and 94% of units pre– and post-MPTP, respectively; Table 1). Positive and negative slopes were equally common among the ramp-like changes detected and that was equally true in both pre– and post-MPTP populations (chi-square test;*χ^2^*(3, n=340) = 0.62, p=0.43; Table 1). Our method for detecting the onset of perimovement changes in activity relative to a SPHP baseline took into account the presence of these ramp-like changes in firing rate.

**Figure 3.**
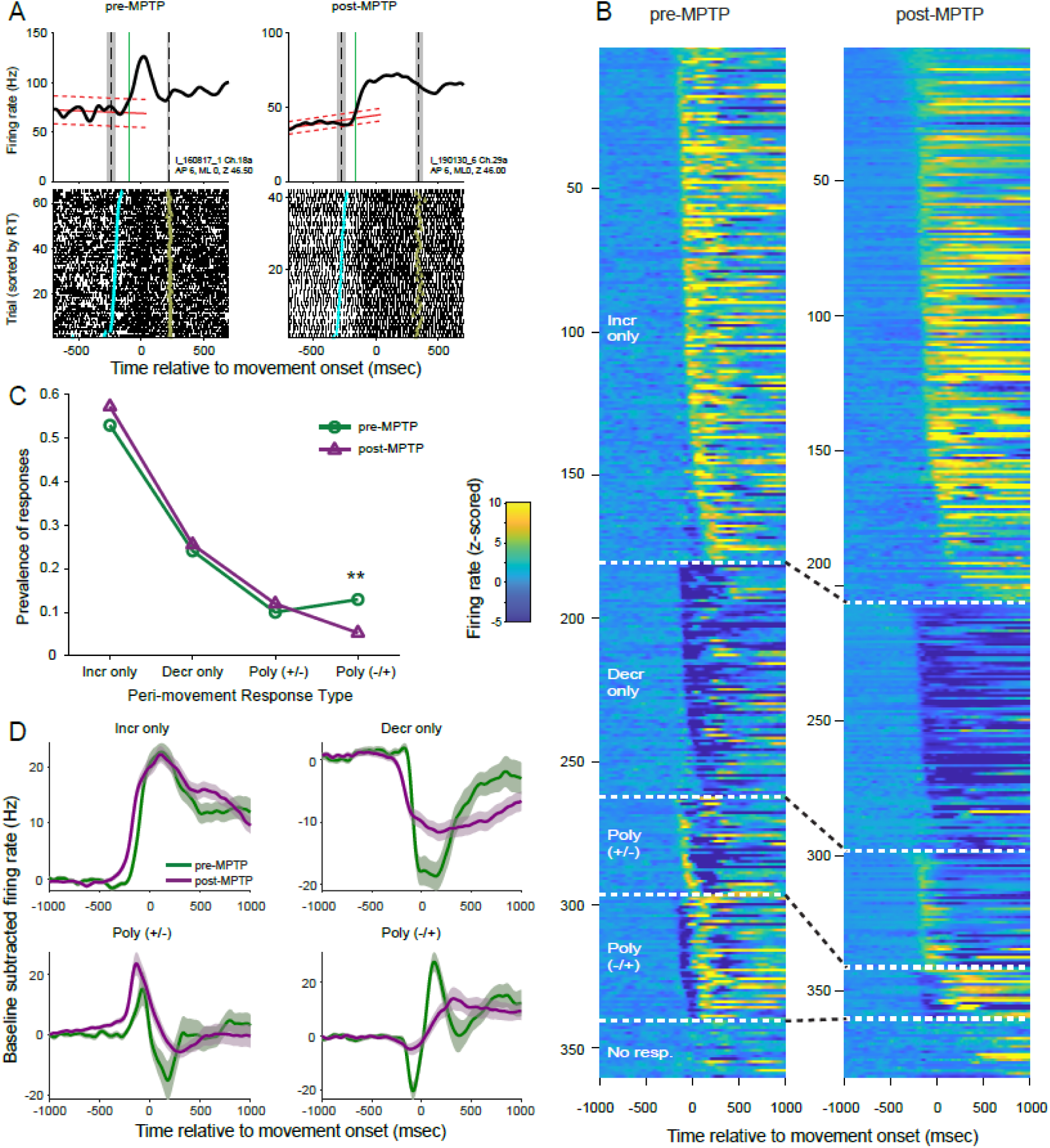
*A*: Peri-movement activity of exemplar GPi single-units sampled pre-(*left*) and post-MPTP (right). Mean SDFs and raster representations of trial-by-trial spike trains are aligned across trials to the time of movement onset (time zero). Vertical black dashed line and gray rectangle indicate the median and IQR of the times of go-cue onset (*left*) and the end of movement (*right*) relative to movement onset. *Sloped red lines*: the unit’s mean baseline trendline ± confidence interval. *Vertical green line*: time of onset of mean response. For raster diagrams, single trial spike trains are sorted according RT, shortest at top. For each trial, colored circles indicate times of go-cue onset (*cyan*) and target contact (*olive*). *B*: Peri-movement SDFs of all single-units studied, sorted according to response type and response onset latency relative to movement onset (earliest onset at the top for each response category). Left and right panels show data from pre– and post-MPTP periods, respectively. SDFs were z-scored relative to the mean baseline activity prior to go-cue onset and displayed on a color scale (*inset*). *C*: Overall proportions of perimovement responses classified into the four response types differed between pre– and post-MPTP conditions (*green* and *magenta*, respectively; p<0.01, chi-square test). The proportion of polyphasic (−/+) responses decreased post-MPTP as compared to the pre-MPTP population (** p<0.01, standardized residual analysis). *D*: Mean population-averaged SDFs constructed separately for each response type and pre– and post-MPTP collection periods (*green* and *magenta*, respectively). *Shaded areas*: ±SEM.

### Perimovement responses persist post-MPTP

Next, we examined how perimovement modulations in firing rate were altered following the induction of parkinsonism. Figure 3A shows the activity of two example single units sampled from nearby sites in the GPi of monkey I, one from the neurologically-normal animal and the other following MPTP administration (left and right subpanels, respectively). Both units showed a monophasic increase in firing that began during the RT interval [*vertical green line* denotes the estimated time of response onset relative to the unit’s baseline activity; i.e., linear trend ±confidence interval (CI) of activity prior to go-cue presentation, *red sloped lines* in Fig. 3A].

Nearly all single-units showed a significant perimovement change in discharge for at least one direction of movement. Contrary to previous report (30), the prevalence of perimovement changes did not increase following MPTP administration (98% of neurons were responsive both pre– and post-MPTP; chi-square test; *χ^2^*(1, n=371) = 0.213, p=0.64).

Perimovement activity differed between movement directions (i.e., was considered directional) in 60% of neurons sampled prior to MPTP administration and that fraction was unchanged following MPTP (51% of neurons post-MPTP; chi-square test; *χ^2^*(1, n=371) = 2.83, p=0.09). That observation was reinforced by results from an alternate approach that tested for directionality of discharge across the whole behavioral response interval (directional discharge in 57% and 54% of neurons pre– and post-MPTP, respectively; chi-square test; *χ^2^*(1, n=371) = 0.18, p=0.67).

The form and timing of neuronal responses differed widely between neurons (Fig. 3B). First, we considered neuronal responses independently for each movement direction and tested for changes in the incidence of different forms of perimovement activity between pre– and post-MPTP populations (Fig. 3C). Monophasic changes in firing (composed of a simple increase or decrease in firing) were the most common, amounting to 79% and 83% of all responses under pre– and post-MPTP conditions, respectively (chi-square test; *χ^2^*(1, n=689) = 2.49, p=0.11). Similarly, the fraction of polyphasic responses was not changed following MPTP (21% pre-MPTP versus 17% post-MPTP). Polyphasic responses were subdivided further according to the sign of the initial change: increase followed by decrease [ *Poly(+/−)*] and decrease followed by increase [ *Poly(−/+)*]. All detected responses are shown sorted by response type and latency in Figure 3B.

Considering all four categories of response type, the prevalence of different response types changed with the induction of parkinsonism (chi-square test; *χ^2^*(3, n=689) = 11.30, p<0.05; Fig. 3C). That effect was attributable to an isolated reduction in the prevalence of Poly(−/+) type responses following MPTP administration (13% of responses pre-MPTP versus 5% post-MPTP; standardized residual analysis; p < 0.01). That reduction was accompanied by slight increases in prevalence of the other three response types following MPTP.

Thus, we found no evidence that parkinsonism was accompanied by a change in the overall prevalence of perimovement modulations in activity. We did find a modest reduction in the incidence of polyphasic responses that began with a decrease in firing.

### Perimovement decreases: preserved prevalence but altered metrics

Previous studies have suggested that parkinsonian motor symptoms arise from a selective loss of GPi perimovement decreases in firing (30, 41). In the current dataset, however, decreases in activity were equally common in the populations collected before and after MPTP administration. As a fraction of all monophasic changes in firing, decreases were equally common in the pre– and post-MPTP populations (31%; chi-square test; *χ^2^*(1, n=558) = 0.004, p=0.95). Furthermore, when the individual phases of all detected responses were considered independently, decreases in firing were also equally common in pre– and post-MPTP populations (chi-square test; *χ^2^*(1, n=820) = 0.105, p=0.75).

Inspection of population averaged SDFs suggests features of responses that changed with the induction of parkinsonism (Fig. 3D). Population responses diverged from baseline firing at earlier latencies preceding movement onset as compared with responses pre-MPTP. That shift to an earlier latency was equally evident for Incr-only, Decr-only and Poly(+/−) response types, but barely perceptible for the Poly(−/+) type. Post-MPTP population responses also tended to have prolonged durations and reduced magnitudes as compared with responses pre-MPTP. Those reductions in response magnitude were most evident for decreases in firing rate (i.e., for the Decr-only response type and selectively for the decrease components of Poly(+/−) and Poly(−/+) response types).

### Perimovement responses are earlier, smaller and longer duration post-MPTP

The results so far suggest that parkinsonism was associated with changes in the timing and shape of perimovement modulations in GPi activity. To gain deeper insight into the specific response features affected, we took independent measurements of response onset latency, response magnitude (peak change from baseline) and response duration (full width at half max, FWHM) for perimovement modulations in the across-trials mean SDFs (see Fig. 4A). For polyphasic responses, those measures were obtained for the initial phase of the response. Measures were then compared between pre– and post-MPTP neural populations. In these analyses, responses for each direction of movement were considered separately.

**Figure 4.**
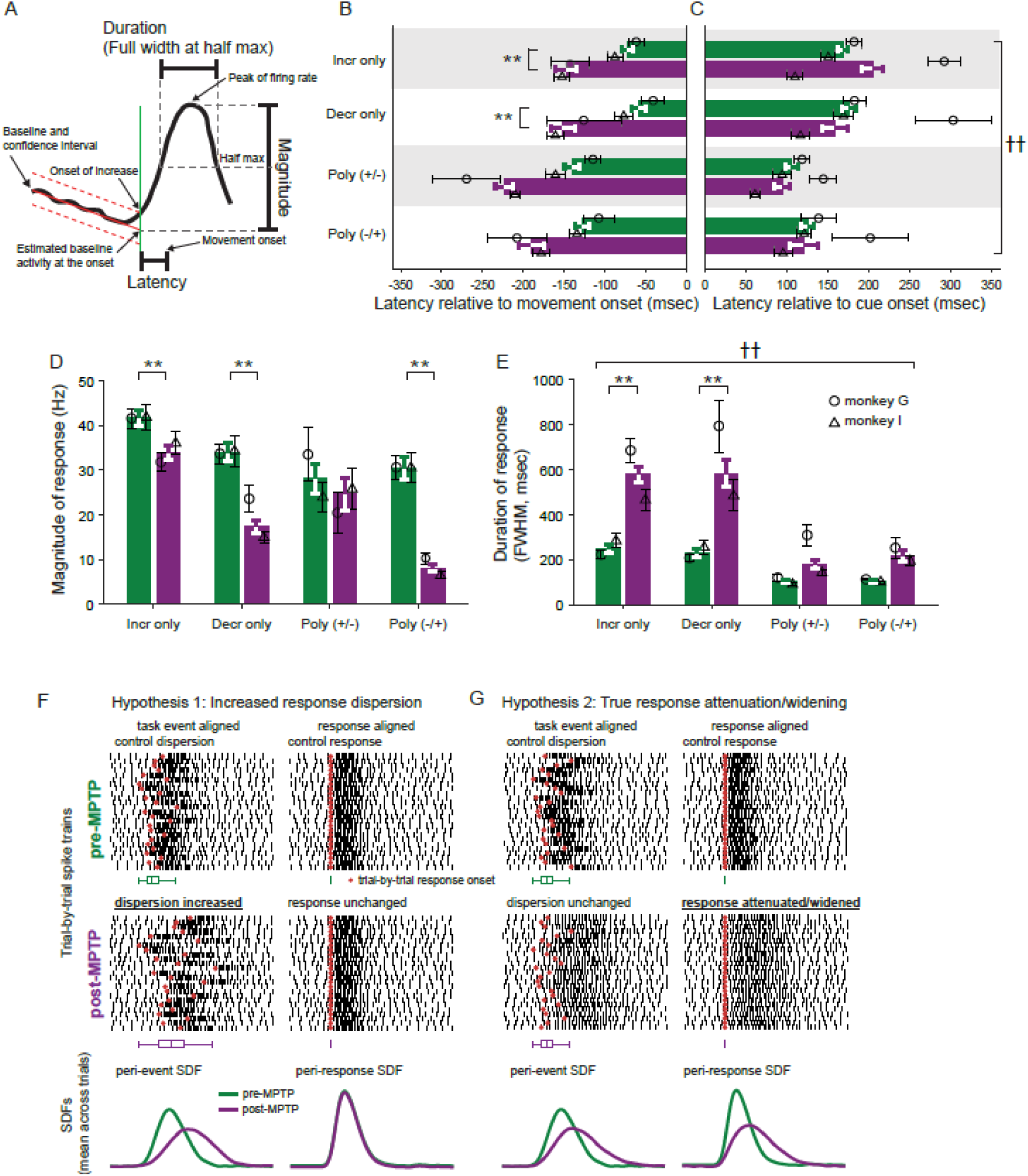
*A*: Response metrics extracted from mean across-trials SDFs included response onset latency, response magnitude, and response duration. (Using conventions from Fig. 3A.) *B-C*: Mean (± SEM) response onset latencies relative to movement onset (*B*) and go-cue (*C*) for each response type (*green*: pre-MPTP, *magenta*: post-MPTP). MPTP administration shifted responses to earlier onset latencies relative to the time of movement onset (*B*: ** p<0.01). Relative to the time of go-cue presentation (*C*), mean latencies did not differ comparing whole pre-versus post-MPTP populations, but latency shifts did differ between the two animals (*B*: ††, p<0.001). Following MPTP, go-cue –to-response latencies were prolonged in animal G (*open circles*) while they were shortened in animal I (*open triangles*). *D*: Mean response magnitudes were attenuated following MPTP administration. That attenuation was more severe for decrease-type responses and the decrease component of Poly(−/+) type responses. *E*: Response durations were prolonged following MPTP administration and that prolongation was accentuated in animal G. *F-G*: Two potential mechanisms underlie the observed response attenuation and prolongation post-MPTP: *F*, an increase in the trial-to-trial temporal dispersion (“jitter”) of responses relative to the task event used for alignment. *G*, a true attenuation and widening of the neuronal response. For both proposed mechanisms, trial-by-trial rasters of simulated spike trains (*top*) and across-trials mean SDFs of those simulated spike trains (*bottom*) are shown as they appear when aligned on a task event (*left*) and aligned on the actual onset of the simulated response (*right*). *Red circles*: times of response onset on individual trials. Below each raster, *horizontal box plots* depict the trial-by-trial distribution of response onsets (IQR).

Following MPTP administration, neuronal responses shifted to earlier onset latencies preceding movement initiation as compared with the onset latencies observed pre-MPTP (3-way ANOVA main effect of MPTP; F(_1,676_)=35.77, p < 0.01; Fig. 4B). Similar-sized shifts to earlier latencies were observed for all response types (3-way ANOVA, MPTP × response-type interaction; F(_3,676_)=0.20, p=0.90) although post-hoc analysis indicated that that shift was significant only for Incr-only and Decr-only response types. In contrast, no significant change in mean latencies was found when response latencies were analyzed relative to the time of go-cue presentation (3-way ANOVA main effect of MPTP; F(_1,652_)=2.20, p = 0.14; Fig. 4C). However, MPTP affected this aspect of response timing differently in the two animals. In animal G, MPTP administration led to a shift toward later onset times following go-cue presentation, with mean latency shifts of +84 ms and +96 ms for increase– and decrease-type responses, respectively (Fig. 4C, circles). In contrast, in animal I, responses shifted to earlier onset times, with mean shifts of −36 ms and −55 ms for increase– and decrease-type responses, respectively (Fig. 4C, triangles; 3-way ANOVA, MPTP × animal interaction (F(_1,652_) = 53.24, p < 0.001). This pattern was consistent across all response types (3-way ANOVA, response type × animal interaction; F(_3,652_) = 0.51, p = 0.67). Notably, the animal that exhibited the prolongation of cue-to-response latencies, monkey G, also displayed the more severe slowing of behavioral measures (RT and MD; see Fig. 1F-G).

Response magnitudes were reduced in size following MPTP (3-way ANOVA, main effect of MPTP; F(_1,669_)=32.52, p<0.01; Fig. 4D). Although some reduction in response magnitude was observed for each response type, the reduction was more severe in decrease-type responses than in increase-type responses (3-way ANOVA, MPTP × response-type interaction; F(_3,669_)=3.61 p<0.05). More specifically, monophasic decreases were reduced in size by 49% whereas monophasic increases were reduced in size by 19%. Similarly, among polyphasic responses, the initial decrease of Poly(−/+) responses saw a 74% reduction in magnitude whereas the initial increase of Poly(+/−) responses declined by only 16%. The effects of MPTP on response magnitudes did not differ significantly between animals (3-way ANOVA effect of MPTP × animal interaction; F(_3,669_)=0.13, p=0.72).

Response durations (FWHM) were prolonged following MPTP (3-way ANOVA, main effect of MPTP; F(_1,638_)=49.38, p<0.01; Fig. 4E). That prolongation was strikingly more severe for monophasic responses (233% and 252% for Incr– and Decr-only response-types, respectively; 3-way ANOVA, MPTP × response-type interaction; F(_3,638_)=3.15, p<0.05) than for polyphasic responses (175% and 211% for Poly(+/−) and Poly(−/+) response-types, respectively). Similar patterns of effects on response magnitude and duration were found when those metrics were extracted from mean SDFs aligned relative to the time of go-cue presentation (Supplemental Fig. S4 A-B).

The alterations in perimovement activity described thus far were taken from across-trials mean SDFs. Those measures, however, are susceptible to distortions that depend on how variable the timing of a response is from trial-to-trial (i.e., its temporal jitter) relative to the task event used to align the across-trials average. Two distinct mechanisms could account for a reduction in mean response magnitude and an increase in mean response duration: 1) an increased dispersion in the trial-to-trial timing of the response, but no change in the true magnitude or duration of the response itself (illustrated in schematic form in Fig. 4F, *Hypothesis 1*); or 2) the response itself is reduced in magnitude and prolonged in duration, while the trial-to-trial jitter in response timing is unchanged (Fig. 4G, *Hypothesis 2*). In other words, when measurements are taken from mean across-trials SDFs, the net effect of an increase in the trial-to-trial jitter of response timing can be indistinguishable from a true reduction in response magnitude and increase in response duration. To address that ambiguity, we sought to control for the trial-to-trial jitter in response timing by: 1) estimating the times of onset of neuronal responses on individual trials; 2) creating response-aligned mean SDFs; and 3) quantifying response magnitude and duration from those response-aligned mean SDFs.

### Trial-by-trial response timing relative to task events

The times of onset of neuronal responses on individual trials were detected using a technique similar to that described previously (62, 63)(see *Materials and Methods*). In addition to its utility for disambiguating response metrics, information on trial-to-trial response timing can provide valuable insights into the temporal (and, potentially, functional) linkages between neuronal responses and behavioral events. Inspection of the perimovement raster plots of many neurons suggested that responses on individual trials began at relatively fixed latencies either following the onset of the go-cue or relative to the time of movement onset (Fig. 5B and C, respectively). Timing relationships like these provide clues into the stage of processing a neural population participates in as part of a sensory-motor RT (e.g., as in the task used here)(62, 77–79). These may include clues into the slowing of the RT process in parkinsonism.

**Figure 5.**
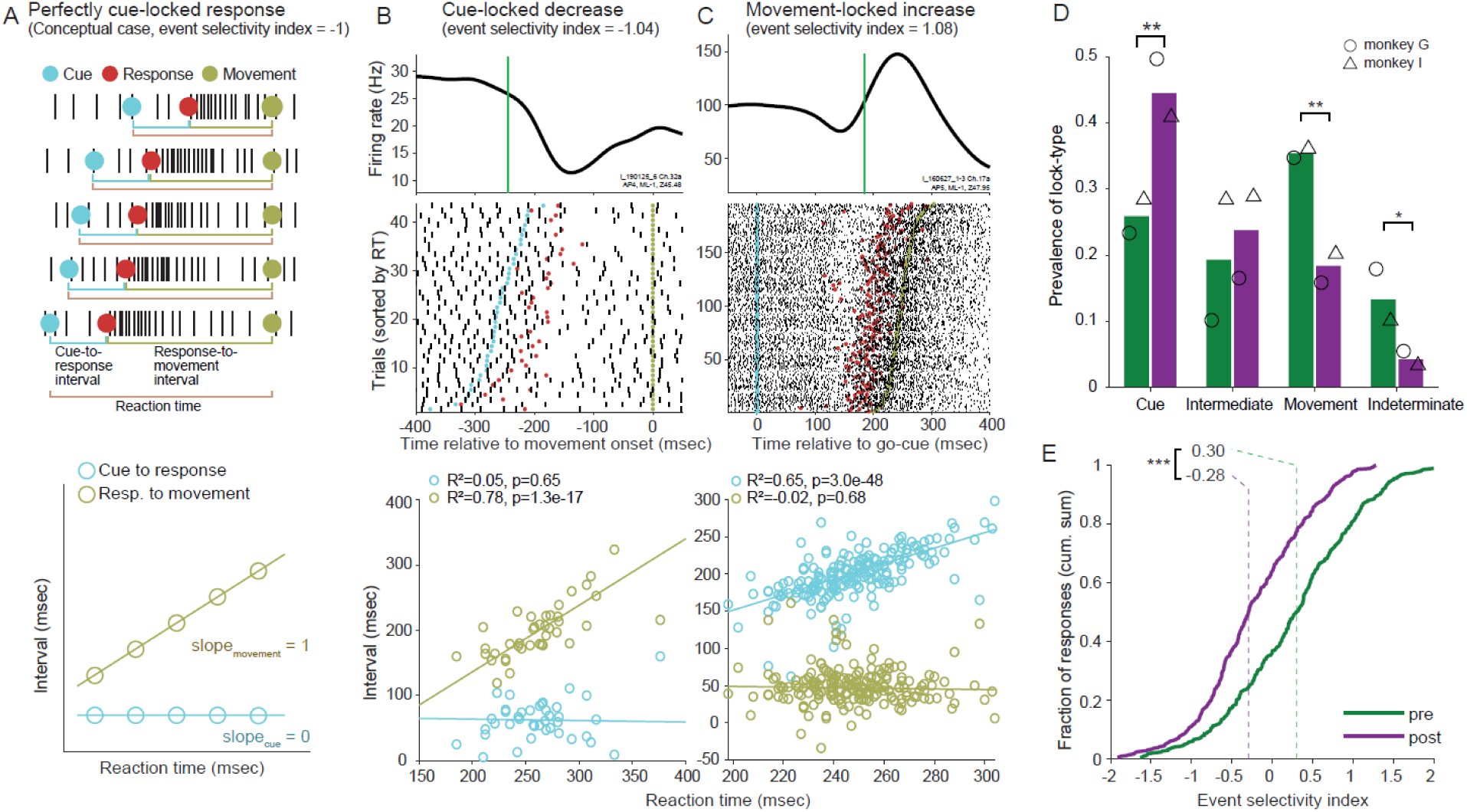
*A*: Event-locking approach. *top*: raster diagram of a simulated neuronal response whose trial-to-trial timing is linked closely to go-cue onset. Spike trains for individual trials are sorted according to RT, shortest RT at top. Time intervals used in the analysis are labeled below. Colored circles show times on individual trials of go-cue onset (*cyan*), neuronal response onset (*red*) and movement onset (*olive*). *bottom*: trial-to-trial values reflecting cue-to-response (*cyan*) and response-to-movement (*olive*) time intervals are plotted relative to the associated single trial RT. Cue-to-response intervals exhibit no relation to the trial-to-trial variations in RTs (i.e., slope = 0), whereas response-to-movement intervals covary strongly with RT (slope = 1). *B-C*: Activity of exemplar GPi units whose responses were categorized as time-locked to the go-cue (*B*) and movement onset (*C*). *B top*: trial-averaged SDF (*black*) and sorted raster (shortest RTs at top). Estimated times of response onset (*red circles*) relative to the times of go-cue (*cyan*) and movement (*olive*) onset. *B bottom*: similar to panel *A*, cue-to-response intervals (*cyan*) exhibit no relation to trial-to-trial variations in RTs, whereas response-to-movement intervals (*olive*) covary strongly with RTs (legend details results from linear regressions). *C*: a response categorized as movement-locked because cue-to-response intervals covaried closely with RTs while response-to-movement intervals did not (following conventions of panel *B*). *D*: Temporal linkage of responses to movement declined post-MPTP. Movement-locked responses were the most common form of event locking prior to MPTP, whereas cue-locked responses became most common post-MPTP. The prevalence of intermediate locking (significant relations for both cue-response and response-movement intervals) did not change following MPTP. *E*: Event selectivity indices (ESIs). ESIs for individual neuronal responses were distributed smoothly ranging from more cue-locked (*negative values*) to more movement-locked (*positive values*). Prior to MPTP, the distribution centered on a median ESI of 0.30, supporting a greater prevalence of locking to movement. Following MPTP, the distribution shifted leftward to favor more cue-locked responses (median –0.28; p<0.001, Kolmogorov-Smirnov test).

Neural responses that participate at different stages of the sensory-motor RT process may be distinguished from each other by the variability in their timing relative to the appearance of the visual go-cue and the onset of movement. If a response has a tight temporal linkage to the sensory go-cue, then the time interval between the go-cue and the neuronal response should be constant regardless of the RT (i.e., slope ≈ 0, see schematic in Fig. 5A), whereas the interval between response onset and movement should covary strongly, across trials, with the behavioral RT (with slope ≈1). If, on the other hand, the response is linked tightly to movement initiation, then the interval between the go-cue and the response should covary with RT (e.g., exemplar single-unit in Fig. 5C). Note that discrimination of those two temporal alignments depends critically on the presence of trial-to-trial variability in the behavioral RT (the interval between go-cue appearance and movement initiation).

To quantify those timing relationships, we calculated two separate linear regressions each comparing the trial-by-trial covariation of a time interval (go-cue–to–response or response–to– movement) with the behavioral RT. A significant positive relation (p<0.05) between response– to–move intervals and RTs, in the absence of a significant relation between cue–to–response intervals and RTs, was taken to indicate that the response was time locked to go-cue onset (i.e., *cue-locked*, Fig. 5A and B). Conversely, a significant positive relation between cue–to–response intervals and RTs, but not between response–to–move intervals and RTs, indicated that the response was time locked to movement onset (*move-locked*; Fig. 5C). If both regressions yielded significant positive slopes, a phenomenon also reported by (79), then the response was categorized as having “intermediate” locking. If neither regression was significant, then the time locking was categorized as *indeterminate*.

Typical examples of single-unit responses time-locked to cue and movement are shown in Figure 5B and C, respectively. Movement-locked responses were the most common form of event locking in the pre-MPTP period (40.9% of responses; chi-square test; *χ^2^*(3, n=677) = 19.0, p<0.01, Fig. 5C), which is consistent with previous observations (62, 63, 80). Cue-locked responses were the second most common (23.0%). Intermediate-type locking (18.9%) and indeterminate event locking (17.2%) were relatively rare.

The prevalence of different event locking types changed markedly following the induction of parkinsonism (chi-square test; *χ^2^*(3, n=820) = 42.70, p<0.001; Fig. 5D). Movement-locking became far less common post-MPTP, dropping to 23.8% of responses, whereas cue-locking increased markedly to become the most common form of event locking (40.8% of responses). The prevalence of intermediate-type locking did not change significantly. Indeterminate locking became very rare in the post-MPTP population. Very similar changes in event locking were found when increase– and decrease-type responses were considered separately (see Supplemental Fig. S5).

To gain deeper insight into the distribution of different forms of event locking, and the interacting effects of MPTP, we calculated an event locking index (*ELI*) that reflects the relative difference in slopes yielded by the two regressions (slope_cue-resp_ and slope_resp-mvt_) as follows: ELI = (slope_cue-resp_ − slope_resp-mvt_)/ (slope_cue-resp_ + slope_resp-mvt_). An ELI value ≤ −1 denotes perfect cue-locking, ELI ≥ +1 denotes perfect movement-locking, and an ELI value of 0 denotes a response whose timing covaries with the midpoint of the RT interval regardless of the overall duration of the RT.

Prior to MPTP administration, ELIs spread across a broad Gaussian-like distribution centered on a median ELI of 0.30 (Fig. 5E). The smooth distribution provided no evidence for the presence of separable categories of event locking. Consistent with that conclusion, clustering analysis indicated that the distribution was most likely composed of one unimodal cluster (see *Methods – Statistics*; (67). These results imply that the discrete categories of event locking described above and previously (62, 63, 80) and Fig. 5D) are a product of the categorical analysis used. The broad smooth distribution of ELIs suggests that GPi responses participate instead at a multitude of steps in the sensory-to-motor processes that occur during the RT interval.

Following the induction of parkinsonism, the overall distribution of ELIs shifted markedly to the left, away from movement-locking and toward more cue-locked responses (median ELI = −0.29; Kolmogorov-Smirnov test; p<1.6 e^-14^; Fig. 5E). The shift was due in part to a selective reduction in responses locked closely to movement onset. Also evident was a trend for the distribution of ELIs to skew further leftward post-MPTP (skew = −0.13 and 5.2 e^-4^ for pre– and post-MPTP, respectively), consistent with an increased representation of responses locked to cue-onset following the induction of parkinsonism.

To summarize, the normally strong temporal linkage of many GPi responses to movement initiation (62, 63, 80) was reduced following the induction of parkinsonism. That was replaced by an increased temporal locking of responses to go-cue onset. That shift in coupling, combined with the prolongation of the response-to-movement interval described above (Fig. 4B), may provide insights into the pathophysiology of increased RTs in parkinsonism (31, 81). Before discussing that idea in more depth, we examine the effects of parkinsonism on the fundamental variability in response timing and compare response metrics after controlling for the trial-to-trial jitter in response timing.

### Effects of MPTP on temporal dispersion of responses differed between animals

The trial-to-trial jitter in timing of neuronal responses became more variable following the induction of parkinsonism in one of the two animals. We measured the residual variability in response timing after regressing out the components of temporal variability attributable to differences in event locking and the trial-to-trial variability in RTs (see *Methods – Analysis* for details).

Considering the mean across both animals, the temporal dispersion of responses increased significantly following MPTP [3-way ANOVA, main effect of MPTP; F(_1,728_)=215.15, p<0.001; IQR (inter-quartile range) in Fig. 6A]. That result, however, was due completely to a marked effect of MPTP in neurons sampled from monkey G (mean 213% and 183% increase in IQR for increase– and decrease-type responses, respectively) whereas minimal changes were found in monkey I (mean 9% decrease and 10% increase in IQR for increase– and decrease-type responses, respectively; 3-way ANOVA, MPTP × animal interaction; F(_1,728_)=224.54, p<0.001). The pattern of effects was very similar for increase– and decrease-type responses (3-way ANOVA, MPTP × response type interaction; F(_1,728_)=0.01, p=0.92) and for responses categorized as cue-locked and movement-locked (see supplemental Fig. S6A and D). It is worth noting that the one animal that showed a large increase in the temporal dispersion of responses, monkey G, was also the animal whose task performance was more severely affected by MPTP (see Fig. 1F-G).

**Figure 6.**
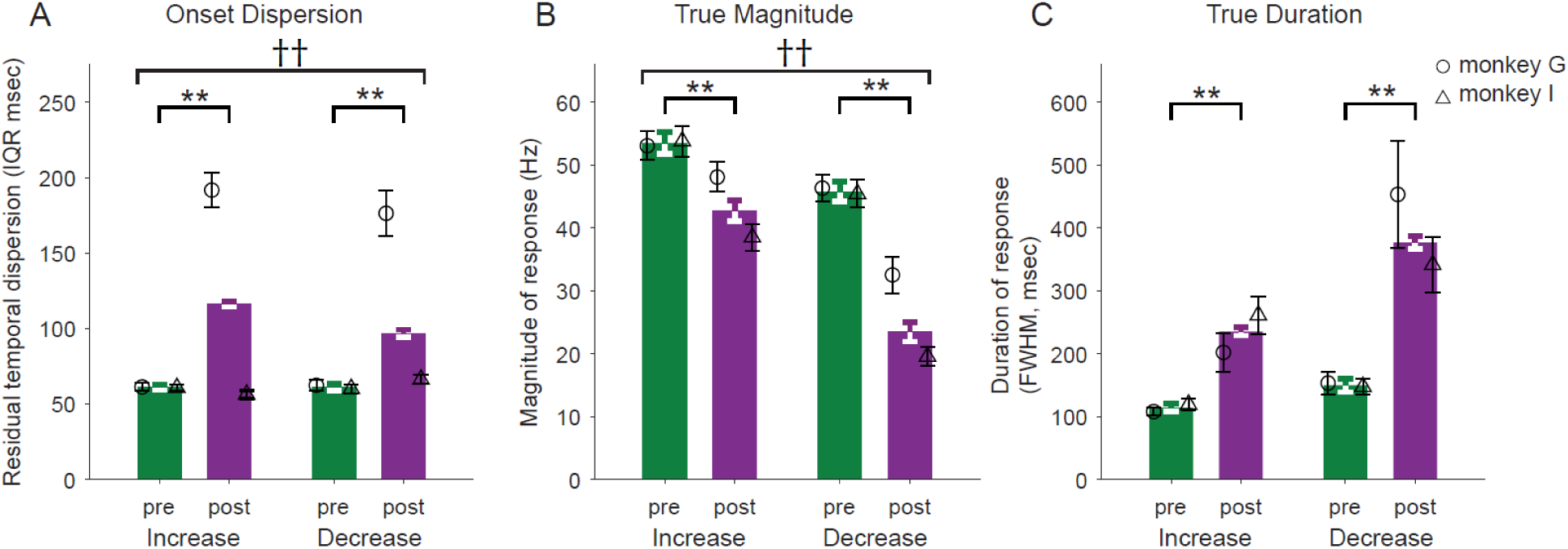
Mean response metrics after controlling for trial-to-trial variability in response timing. *A*: Temporal dispersion of responses after controlling for time-locking to events. *B*. Response magnitude from response-aligned spike-density function. *C*: Response duration from response-aligned spike-density function. Pre– and post-MPTP (*green* and *magenta*, respectively). Statistical differences computed using 3-way ANOVA followed by Tukey’s test (** p<0.01 main effects of MPTP; †† p<0.01 animal-by-MPTP interaction). All values are presented as mean ± SEM. Open circles: monkey G. Open triangles: monkey I.

### De-jittered responses confirm smaller and wider responses post-MPTP

We suggested above that the MPTP-induced alterations in trial-averaged responses (i.e., reduced magnitude and prolonged duration; Fig. 4D, E) might be attributable to an underlying increase in the temporal jitter of response onset times across trials. That idea can now be addressed by using the trial-by-trial response onsets to re-align single trial SDFs (thereby removing the effects of temporal jitter; see *Methods – Analysis*), averaging across trials and then extracting response metrics from the response-aligned SDFs. Polyphasic response types were excluded from this analysis due to the relatively small number of responses of those types.

Metrics taken from de-jittered responses showed effects that were relatively similar to those extracted from trial-averaged SDFs. De-jittered responses showed a reduction in magnitude following MPTP (3-way ANOVA main effect of MPTP; F(_1,703_)=76.46, p<0.001; Fig. 6B). Although both increase and decrease response types showed significant reductions, decrease-type responses showed a stronger reduction in magnitude than increase-type responses (3-way ANOVA MPTP × response-type interaction; F(_1,703_)=6.86, p<0.01). Following the induction of parkinsonism, monophasic decreases were reduced in size by 48% whereas monophasic increases were reduced by 21%. The effect of MPTP on the de-jittered response magnitude was significantly different between the two animals (3-way ANOVA MPTP × animal interaction; F(_1,703_)=9.85, p<0.01). The animal that showed the larger decrease in response magnitude, monkey I, was the animal whose behavior was *less* severely affected by MPTP. We will discuss this contradictory observation below.

Similarly, the durations of de-jittered responses were prolonged following MPTP (3-way ANOVA main effect of MPTP; F(_1,657_)=64.34, p<0.001; Fig. 6C). Notably, that prolongation was greater for decrease-type responses than for increase-type responses (106% and 152% for increase– and decrease-type responses, respectively; 3-way ANOVA MPTP × response-type interaction; F(_3,657_)=6.33, p<0.05). Note that this extra prolongation of decrease-type responses was not seen in metrics taken from trial-averaged SDFs (Fig. 4E) and thus was uncovered by the de-jittering process.

We performed an additional analysis to investigate potential influences on these results of the MPTP-induced changes in baseline firing rates (detailed above and in Table 1). We normalized individual response-aligned SDFs using a standard Z-scoring method based on trial-to-trial firing rates from the baseline period. Comparison of response magnitudes and durations extracted from these normalized SDFs yielded results that were similar in most respects to those described above. The magnitudes of decrease-type responses were smaller post-MPTP and durations of both types of responses were prolonged (Supplemental Fig. S7A and B, respectively; ANOVA results in figure). Increase-type responses, however, appeared to increase in magnitude post-MPTP after normalizing for firing rate. This result further highlights the differing effects of MPTP-induced parkinsonism on increase– and decrease-type GPi responses. The potential significance of an exaggeration in increase-type responses is discussed below.

In conclusion, even after controlling for changes in the trial-to-trial jitter of response timing and changes in baseline firing rates, the magnitudes of decrease-type responses were diminished and response durations were prolonged following the induction of parkinsonism.

## DISCUSSION

We investigated the effects of MPTP-induced parkinsonism on the perimovement activity of neurons in the primate GPi. We found that parkinsonism was associated with a shift in the timing of perimovement responses to earlier onset times, reduced temporal linkage to movement and a weakening and prolongation of those changes. A particularly noteworthy result was that the weakening and prolongation of responses was more dramatic for decreases in firing as compared with those for increase-type responses. These results confirm and expand upon a previous description of parkinsonism-related abnormalities in GPi movement-related activity (30). Given that GPi neurons form the principal efferent pathway by which the BG influences motor control centers, these changes in GPi perimovement activity may be important factors in the pathophysiology of the motor symptoms of PD.

### Baseline activity in GPi

The mean firing rate of GPi neurons during periods of attentive rest was reduced following the induction of parkinsonism (Table 1). That result conflicts with predictions of the classical pathophysiologic model for PD (8, 9) and the many empirical observations consistent with that model (13, 30, 70, 82–84). However, we are by no means the first to find changes in resting activity that are at odds with the classical model (41, 57, 73, 85, 86) (12, 87–92). How to explain these divergent findings is unclear at present. Ex vivo electrophysiologic studies of SNr neurons in rodent models reported decreased membrane excitability (87) and altered synaptic properties (87, 88, 93, 94) following dopamine depletion. These factors and changes in the spontaneous activity of synaptic inputs from upstream areas may have interacting affects on the firing rate of BG output neurons at rest. The degree of these effects may differ between studies as a function of the severity of parkinsonism induced and the specific intoxication protocol used. Although less likely, the estimate of resting firing rates may also be influenced by differences between studies in the techniques used for recording or signal processing. An additional potential explanation that applies to the present study in particular is that we measured mean firing rates while animals were engaged in a well-learned, attention-demanding behavioral task. Most previous studies recorded from animals that were not performing a defined behavioral task and, often in those studies, the animal’s behavioral state was not well controlled. Here, by measuring mean firing rates during the periods of attentive rest defined by the task’s start position hold intervals, we compared GPi activity during the same relatively well-controlled behavioral state before and after the induction of parkinsonism. The somewhat unexpected effect on mean firing rates found here may be related to the fact that our animals were engaged in a learned behavioral task combined with the possibility that the effects of parkinsonism on mean GPi firing rates differ between behavioral states. Engagement-dependent change in neuronal activity have been studied extensively in healthy human subjects (95)(and references therein). The degree to which those changes extend to the basal ganglia and in parkinsonism remains to be determined.

In other respects, our results were consistent with previous descriptions of parkinsonism-related changes in GPi resting activity (Table 1). The variability in inter-spike intervals and the mean proportion of spikes in bursts increased following the induction of parkinsonism, consistent with previous studies (57, 69, 70, 82, 96). In addition, the fraction of neurons with rhythmic modulations in firing in the beta frequency range increased with the induction of parkinsonism and the prevalence of gamma band oscillations decreased, in agreement with many previous studies (71, 72). Overall, the changes in GPi baseline activity found here were consistent with the animals’ parkinsonian status following MPTP administration.

### Preservation of perimovement activity post-MPTP

In many ways, the perimovement activity of GPi neurons was relatively unaffected by the induction of parkinsonism. The overall fraction of single units showing significant modulations in perimovement activity was high both before and following MPTP administration. Furthermore, the fundamental shapes of the responses detected and the rates at which those shapes appeared were roughly similar between pre– and post-MPTP conditions (Fig. 3C). Only the polyphasic(−/+) type, which accounted for a small fraction of responses pre-MPTP, became even less common following MPTP. These results differ from those of Leblois et al. (30) who reported that an increased fraction of neurons showed perimovement responses following the induction of parkinsonism (49% and 67% pre– and post-MPTP, respectively). Leblois et al. also reported a dramatic decrease in the fraction of neurons showing perimovement decreases in firing (from 54% of neurons pre-MPTP to only 7% of neurons post-MPTP). Again, the reason for these differences between studies is uncertain, but may be attributable to differences in the animal model, severity of symptoms, or the methods used. Worth noting, however, is that we did find a differential diminution in the magnitude of decrease-type responses with the induction of parkinsonism, a result that will be discussed in detail below.

### Timing of GPi responses is uncoupled from movement initiation post-MPTP

Following the induction of parkinsonism, perimovement responses shifted to earlier mean onset times relative to movement initiation (Fig. 4B). In contrast, the effect of parkinsonism on the latencies from go-cue delivery to mean response onset differed between the two animals. A similar pattern of results was described by Leblois et al. (30), including the specific inconsistency in results between animals. In addition, we found that the normally-common trial-to-trial linkage of GPi responses to movement onset was markedly reduced following the induction of parkinsonism (Fig. 5D-E). This second finding has not, to our knowledge, been describe previously.

Those two results provide new clues into the neural mechanisms that underpin the prolongation of the RT interval in parkinsonism (31–33). Consistent with the animals’ parkinsonian status, we found that mean RTs increased significantly – markedly so in one animal – and the trial-to-trial variability in RTs increased as well (Fig. 1F). For each of these two effects on RTs, we identified a respective neural correlate: 1) the change in mean response latency relative to go-cue and movement and 2) the change in trial-to-trial locking relative to go-cue and movement. For both neural correlates, the change induced by parkinsonism was consistent with a breakdown in the temporal coupling of GPi responses to movement onset.

The observation that latencies between go-cue delivery and response onset were affected modestly and inconsistently suggests that, in circuits upstream of the GPi, the *timing* of signal propagation is relatively unchanged following the induction of parkinsonism. That view is consistent with a recent study of the effects of parkinsonism on the propagation of signals evoked by cortical stimulation via cortico-BG direct, indirect and hyperdirect pathways (41). In that study, parkinsonism strongly affected the magnitudes of signals propagated via the three pathways, but the timing of those signals was unaffected. These results suggest that slowing in the conduction of neuronal signals, which is hypothesized to give rise to parkinsonian bradykinesia (34, 35, 97), is not present uniformly across all brain circuits. Moreover and perhaps surprisingly, slowed conduction is not apparent in the circuits immediately upstream of the GPi – circuits known to be affected in multiple ways by the loss of dopaminergic innervation (98, 99).

We found that the time interval between neural response onset and movement was both prolonged and more variable between trials following the administration of MPTP. One conceptually simple mechanism that could account for those effects is the presence, in circuits downstream of the GPi, of some form of pathology that causes a slowed and more variable conduction of the neural responses transmitted from the GPi. There is little evidence for frank pathology in the GPi itself in PD (100), aside from the loss of dopaminergic innervation (101, 102). A variety of disease-related alterations have been described for the thalamic, midbrain, and brainstem areas that receive monosynaptic projections from GPi (23, 103–108). Pathologic changes have also been described for motor cortical areas, which are positioned two synapses downstream of GPi (109–111). Some combination of these pathologic changes may slow the downstream transmission of GPi task-related responses and, thereby, cause a prolongation of RTs. Of course this model assumes that responses transmitted from the GPi play a central role in movement initiation – an assumption that remains open to debate (40, 112–114).

An alternate idea is that abnormalities in GPi perimovement activity may play an active role in disrupting and slowing normal movement initiation. That idea is explored in more detail in the next section. A third possibility is that the prolongation of RTs in parkinsonism is attributable to slowed conduction along neural processing pathways that run in parallel to the GPi but do not involve it directly.

Regardless of the underlying physiologic mechanism, our results suggest that the proximate cause for RT slowing lies either downstream of the GPi or along an independent parallel processing pathway. Recordings from regions directly downstream of the GPi should provide additional clarification regarding these potential mechanisms.

### Response magnitudes, especially of decreases, are attenuated post-MPTP

Perimovement changes in GPi activity were diminished in size and prolonged in duration following the induction of parkinsonism. Although reductions in magnitude were observed in both increase– and decrease-types of responses, perimovement decreases in firing were affected more severely. Monophasic decreases were reduced in magnitude by almost 50% whereas monophasic increases were reduced by 19%, on average. Alternatively, when results were normalized to take into account differences in baseline firing rates, increase-type responses actually increased in magnitude post-MPTP. Those effects persisted after controlling for the possibility that the observed attenuation and prolongation of mean responses could be explained by increases in the trial-to-trial jitter of response timing.

The results found here expand upon those from Leblois et al. (30) yet differ in detail. Leblois et al. report that perimovement decreases in firing became rare following the induction of parkinsonism. Decrease-type responses to proprioceptive perturbations showed a similar selective reduction in occurrence (30, 115). We did not observe a reduction in the rate of occurrence of decrease-type responses, but did find that perimovement decreases were attenuated in magnitude preferentially in comparison with the effects on perimovement increases. This disparity in results may be explained by differences between studies in the severity of symptoms, the task used or the analysis methods applied. Most obviously, what may be detected as a reduction in response magnitude in one study could easily appear as a reduction in occurrence in another depending on the study’s statistical power and analysis methods applied. Our results agree, however, with Leblois et al.’s overall conclusion that parkinsonism is associated with a differential attenuation of perimovement decreases in firing. Generalizing beyond neural correlates of active movement, Chiken et al. (41) studied the effects of parkinsonism on responses evoked in GPi by electrical stimulation of sites in cortex. They found a marked attenuation of decrease-type responses – responses thought to be mediated via conduction of signals along the pro-kinetic cortex◊striatum◊GPi “direct” pathway (40). The direct pathway originates in striatal projection neurons that express the D1 dopamine receptor (D1-SPNs). Excitation of normally-functioning D1-SPNs produces a monosynaptic GABA-mediated inhibition of GPi neurons and thereby, transient decreases in GPi firing. Thus, a degradation of D1-SPN function would lead to a differential attenuation of GPi decreases in firing.

Several lines of evidence support the view that function of the direct pathway is selectively impaired in parkinsonism. D1-SPNs show marked reductions in spine density and dendritic complexity following chronic dopaminergic denervation (116, 117). Multiple studies report that dopamine denervation reduces the intrinsic excitability of D1-SPNs (reviewed in 118) although not all studies concur (116, 119). In addition, the inhibitory influence of D1-SPNs input to BG output neurons is weakened following dopamine depletion (88). Finally, selective excitation of D1-SPNs restores motor function in a rodent model of PD (120) mediated by an apparent increase in the magnitude of pauses in firing of BG output neurons (121).

The canonical model of BG function states that, in the dopamine-replete brain, task-related reductions in GPi activity facilitate movement through disinhibition of neurons in BG-recipient regions of motor thalamus and midbrain (122, 123). Thus, in parkinsonism, a selective attenuation of decrease-type responses, as seen here, would result in deficient perimovement disinhibition of motor circuits and thereby hypokinetic/bradykinetic movements. According to this model, the slowing of movement seen in parkinsonism can be attributed to an attenuation of task-related decreases in GPi firing and their disinhibitory effect on downstream motor circuits.

An alternative model arises from the longstanding and well-supported idea that the BG plays an central role in procedural learning (124–126). For example, Yttri & Dudman (127) showed that selective excitation of D1-SPNs, which is known to increase the magnitude of pauses in BG output neurons (121), results in a persistent bias toward expressing the behavior that was performed during that excitation. Those results suggest that decrease-type changes in GPi activity serve as a reinforcement learning signal in downstream brain areas. Thus in parkinsonism, a persistent attenuation of GPi task-related decreases, as observed here, may lead to impairments in motor performance through a process of repeated inadequate reinforcement, which could also be termed aberrant learning. Further credence for this alternate model of PD pathophysiology comes from the observation that parkinsonian impairments on motor tasks worsen the more the motor task is practiced (128–130).

The magnitude of task-related increases in firing was also altered following MPTP, but the direction of the change depended on whether results were normalized according baseline firing rates. After normalizing for baseline firing rates, increase-type responses were strengthened following the induction of parkinsonism. The two most prominent sources of net excitatory drive to GPi neurons are the glutamatergic inputs from the subthalamic nucleus and facilitation-through-disinhibition effects from the external pallidum. MPTP-induced changes in the magnitude of task-related increases may be attributable to pathologic changes that have been described for both pathways (131–135). Chiken et al. (41) report that the effect of parkinsonism on stimulation-evoked increases in GPi activity differed depending on what temporal component of the response is measured – short latency responses increased in magnitude whereas long latency increase-type components were unchanged. Those results buttress the general view that the change observed in increase-type responses in the parkinsonian state depends on how they are quantified. The functional consequences of these effects remain a matter of speculation.

These results speak to a longstanding debate in parkinsonian pathophysiology on whether hypokinesis is due to an insufficient scaling of motor commands versus an insufficient focusing of action selection (36, 97, 123, 136, 137). Our observation of a selective weakening of task-related decreases in GPi activity, which according to this model are pro-kinetic, is in line with predictions of the scaling hypothesis. The effects observed in increase-type responses were ambiguous and thus difficult to draw conclusions from. In contrast, several aspects of our results are inconsistent with the focusing hypothesis. We found little evidence for a decrease in the specificity of GPi responses, as is predicted by the focusing hypothesis. We found no change in the overall prevalence of task-related responses with the induction of parkinsonism (Table 1) and only a slight shift in the distribution of different response types (Fig. 3C). In addition, the fraction of neurons with responses selective for the direction of movement did not decrease following MPTP administration, as would be predicted by the focusing hypothesis. Thus, in balance, our results are more consistent with the scaling hypothesis.

An important caveat, however, is that the MPTP-induced attenuation of response magnitude did not covary with between-animal differences in symptom severity. The animal that showed greater response attenuation, monkey I, was the animal whose task performance was *less* severely affected by MPTP (see Fig. 6B). Thus, reductions in response magnitude may be associated with parkinsonism without being direct contributors to the expression of motor symptoms.

### Responses durations, especially of decreases, are prolonged post-MPTP

Perimovement changes in activity were also prolonged in duration following the induction of parkinsonism. Similar to the observed effects on response magnitude, the durations of decrease-type responses were prolonged to a greater degree than those of increase-type responses (Fig. 6C; after controlling for the confounding effects of increased trial-to-trial jitter). This similarity suggests that the two effects, depressed response magnitude and prolonged duration, arise from similar pathophysiologic mechanisms. What those mechanisms are remains a matter of speculation, but may be related to some combination of the pathologic changes described above for direct, indirect, and hyperdirect pathways (116–118, 131–135). The consequences of prolonged response duration and relation to pathophysiology are also uncertain.

Independent of the applicability of the classical firing rate model, and the scaling and focusing hypotheses mentioned above, abnormalities in the timing, magnitude and duration of perimovement responses may disrupt the normal function of downstream motor control circuits and thereby play a direct role in the expression of parkinsonian symptoms. To elaborate, abnormalities in the task-related responses transmitted from the parkinsonian GPi may impair the ability of downstream motor control circuits to play-out motor commands in a timely and orderly fashion. In that way, abnormal GPi responses might give rise to slowed initiation and execution of movement (i.e., prolonged RTs and MDs). Studies of the effects of parkinsonism on perimovement activity in regions downstream of the GPi will provide a way to test that possibility (14–17, 138).

### Limitations

One inherent limitation of the present work is that the results are correlative. We cannot be certain if the observed abnormalities in GPi responses cause changes in behavior, or reflect those changes or are merely correlative. The fact that most GPi responses began during the RT interval places GPi responses at an appropriate timing to influence the initiation and execution of movement, but a causal role is by no means guaranteed. Experiments that perturb circuit function will be of assistance in clarifying the causal roles of the abnormalities described here.

A second caveat is that some of the MPTP-induced changes in response metrics differed significantly between animals. A particularly striking example was the attenuation of response magnitudes, which was more severe in the animal whose behavior was *less* severely affected by MPTP (monkey I, Fig. 6B). That result suggests that relationships between altered neuronal response metrics and symptom severity may be non-linear. Similar markedly non-linear relationships have been observed in the relationship between abnormalities in the resting activity of BG output neurons and the degree of disease progression in mouse models of parkinsonism (86). Better understanding of the relationship between GPi task-related responses and parkinsonian symptoms depends on a systematic examination of the abnormalities in neural activity across incremental stages of parkinsonism.

Third, it is important to acknowledge the simplistic nature of the concept, used widely here, of circuits “upstream” and “downstream” of the GPi. BG-thalamocortical circuits are organized as re-entrant loop circuits in which signals transmitted from the GPi can influence the very brain structures that send signals into that BG circuit (139). Considering the typical transit times for these circuits (41, 42) and the durations of GPi responses, there is ample time during a GPi response for abnormal GPi output to loop back and influence the function of the “upstream” circuits that are generating that response. This re-entrant circuit organization may make it possible for the effects of even subtle dysfunctions of the circuit to be amplified as signals are passed several times through the circuit (140).

### Conclusions

In summary, remarkably few studies have examined the effects of parkinsonism on the perimovement activity of neurons in the GPi, the principal output nucleus of the primate BG. We found that perimovement responses were present in the parkinsonian brain at roughly the same overall abundance and distribution of different response types as seen in the neurologically normal state. Nonetheless, parkinsonism was associated with three principal abnormalities in perimovement activity: 1) The timing of GPi responses became uncoupled from movement onset with respect to both mean latency and trial-to-trial variability in timing. 2) Response magnitudes were attenuated. 3) Response durations were prolonged. The effects on both response magnitude and duration were accentuated in decrease-type responses. These abnormalities in GPi perimovement responses may contribute to the pathophysiology of parkinsonian motor symptoms. The differential attenuation of decrease-type responses is consistent with the concept that decrease-type GPi responses are pro-kinetic and that their attenuation may play a role in the pathophysiology of parkinsonism. The potential significance of that observation is tempered, however, by the inconsistent relationship of attenuated response magnitude to symptom severity between the two animals. Future studies at additional levels of symptom severity are likely to clarify the pathophysiologic significance of the changes described here. Also needed are studies of the effects of parkinsonism on perimovement activity in brain regions immediately downstream of the GPi.

## SUPPLEMENTAL MATERIALS

Supplementary Figs. S1-S7 DOI: 10.5281/zenodo.15538407

## DATA AVAILABILITY

Both electrophysiological data and behavioral data are available on the DANDI Archive at DANDI:000947/0.240510.2211. All MatLab code and the processed source data sufficient to reproduce all figures and tables are in preparation and will be available on Zenodo (DOI: 10.5281/zenodo.15276457).

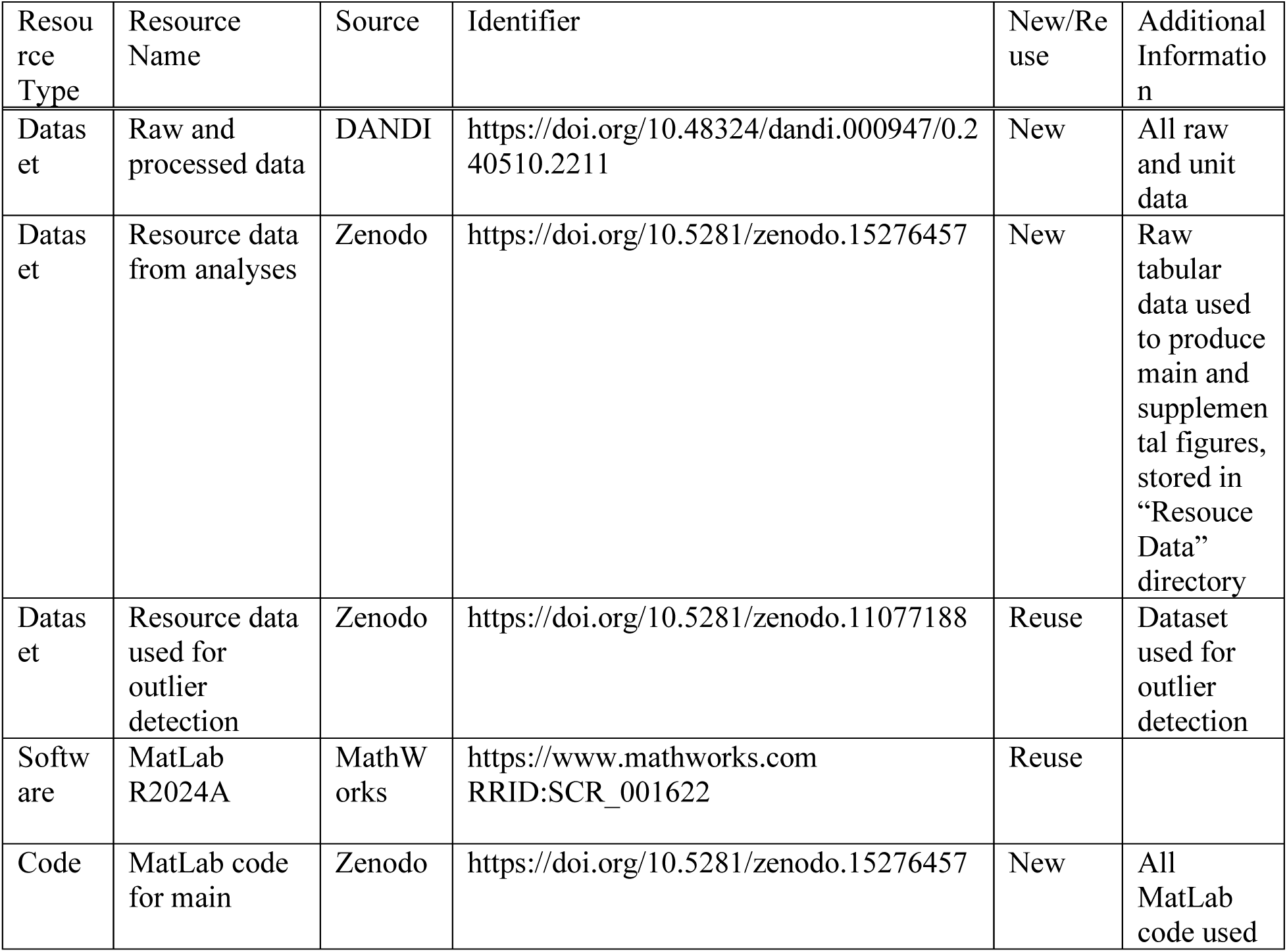
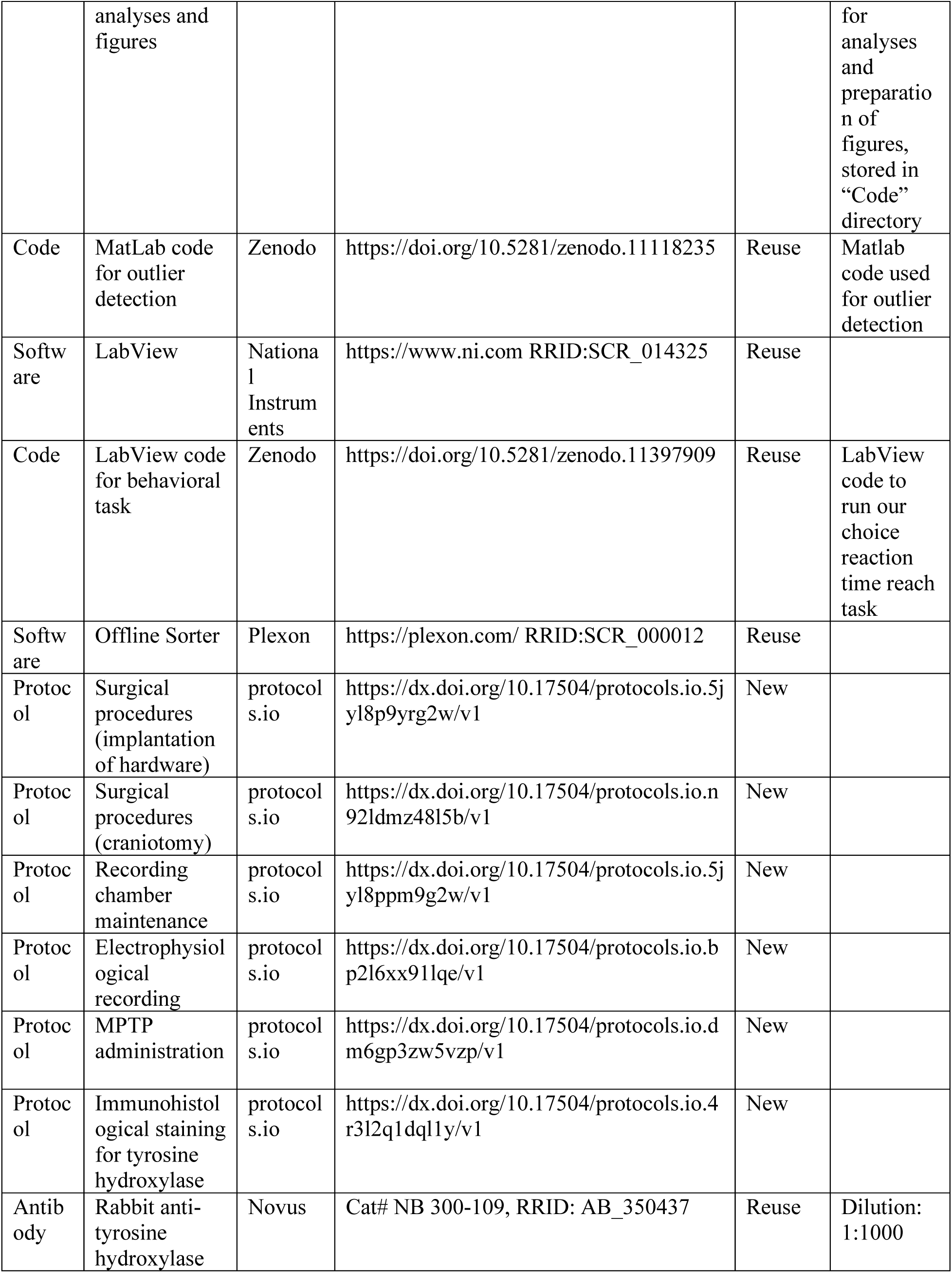
KEY RESOURCE TABLE.

## AUTHOR CONTRIBUTIONS

RST conceived and designed research; DK, AJZ, YH, ACB,, and RMR performed experiments; DK, AJZ, DRH, SB, and KMC analyzed data; DK and RST interpreted results of experiments; DK and RST prepared figures; DK, KMC, and RST drafted manuscript; DK, AJZ, YH, DRH, SB, KMC, ACB, RMR, and RST edited and revised manuscript and approved final version of manuscript.

## ACKNOWLEDGMENTS

We thank Lisa Nieman-Vento and Cherie Lee Cornmesser for their contributions to animal care.

This work was supported by the National Institute of Neurological Disorders and Stroke at the National Institutes of Health (Grant Numbers R01 NS117058-01 and R01 NS070865-01A1 to RST; www.ninds.nih.gov/). Additionally, this research was funded in part by Aligning Science Across Parkinson’s (ASAP-020519) through the Michael J. Fox Foundation for Parkinson’s Research (MJFF). For the purpose of open access, the author has applied a CC BY public copyright license to all Author Accepted Manuscripts arising from this submission.

## SUPPLEMENTAL FIGURE

**Figure S1.**
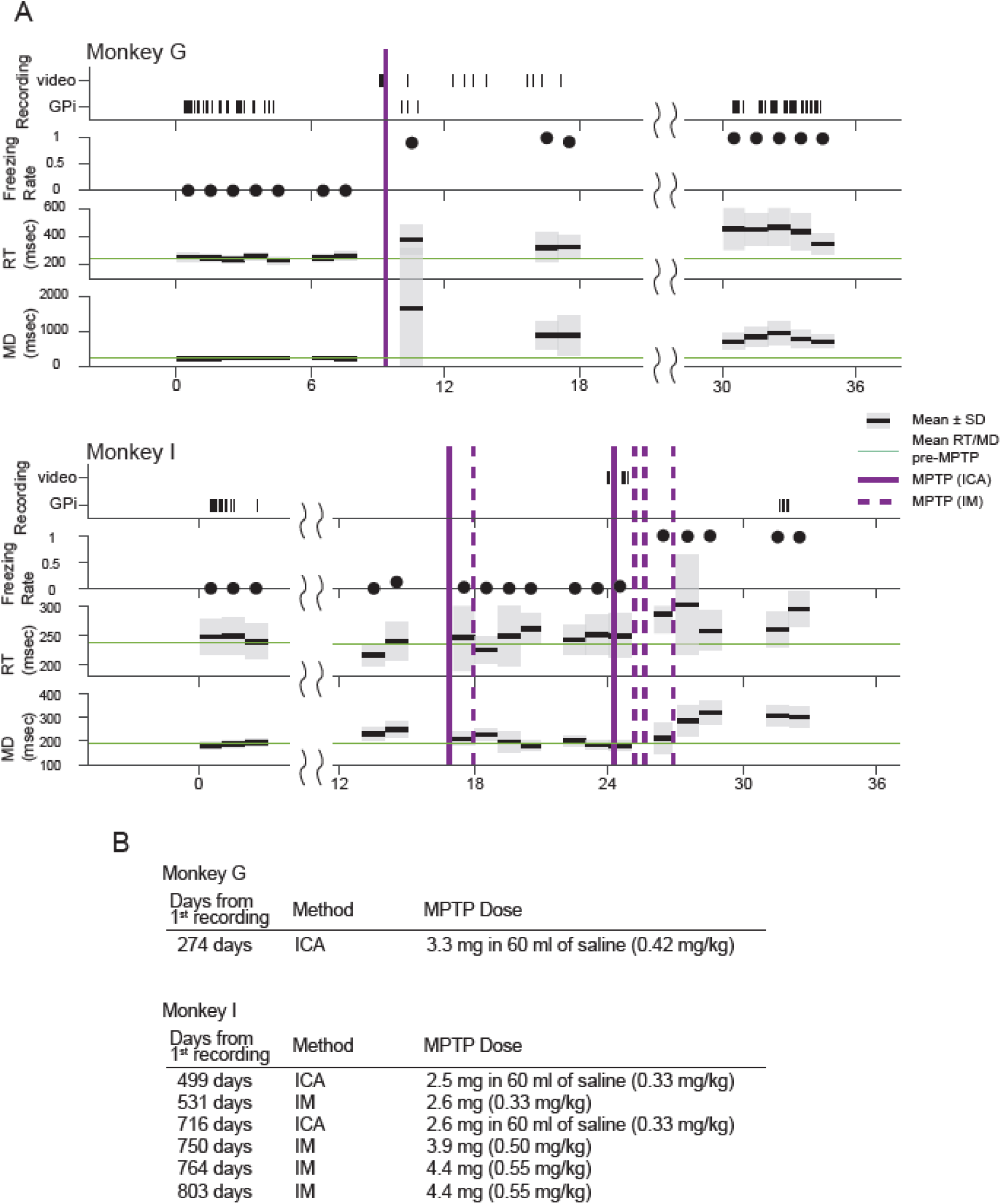
Summary of the experimental timeline. The top and bottom panels correspond to monkeys G and I, respectively. In each panel, tick marks indicate the dates of recording sessions – video recordings of behavior in the home cage (*top of the top row*) and recordings of extracellular neural activity used in the current study (*bottom of the top row*). The rate of occurrence of freezing-like behaviors, reaction times, and movement durations are shown in respective rows second from top to bottom and plotted as means for each month data were collected. *Shading*: +/−SD for reaction time and movement duration. In reaction time and movement duration panels *horizontal green lines*: mean values across pre-MPTP sessions. *Vertical magenta lines*: times of MPTP administration via intra-carotid (*solid lines*) and IM (*dashed lines*) routes. Additional details on MPTP administration are provided below the timeline.

**Figure S2.**
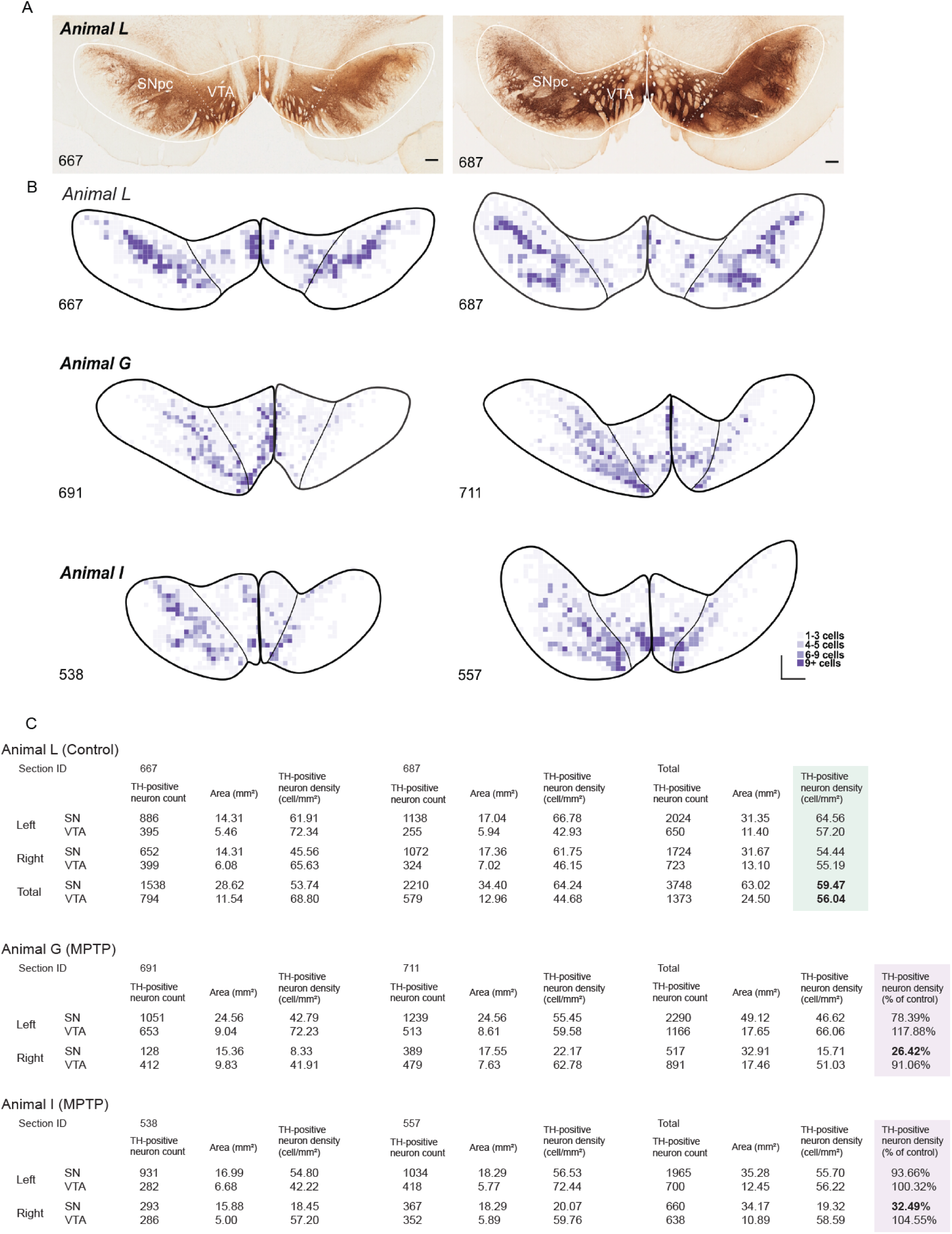
Tyrosine hydroxylase (TH)-positive cells were depleted preferentially from the right substantia nigra (SN) following MPTP administration. *A*. Immunohistochemical labeling for TH in the healthy control animal L revealed dense populations of TH-positive somata in both the SN and ventral tegmental area (VTA). *B*. Density maps of TH-positive somata in the SN and VTA demonstrate a marked depletion from the right SN of animal G (middle) and I (bottom), with no such depletion observed in the healthy control animal L (top). Outlines show estimated borders between VTA and SNpc for two 50 um think sections spaced 1 mm apart through the midbrain of animals L, G and I. Color squares indicate the number of infected neurons in successive 200-um x 200-um bins (see color key). *C*. Summary of the number and density of TH-positive somata identified in the SN and VTA. TH-positive neurons were depleted markedly yet selectively from the right SN (bold text in far-right column) in both MPTP-treated animals.

**Figure S3.**
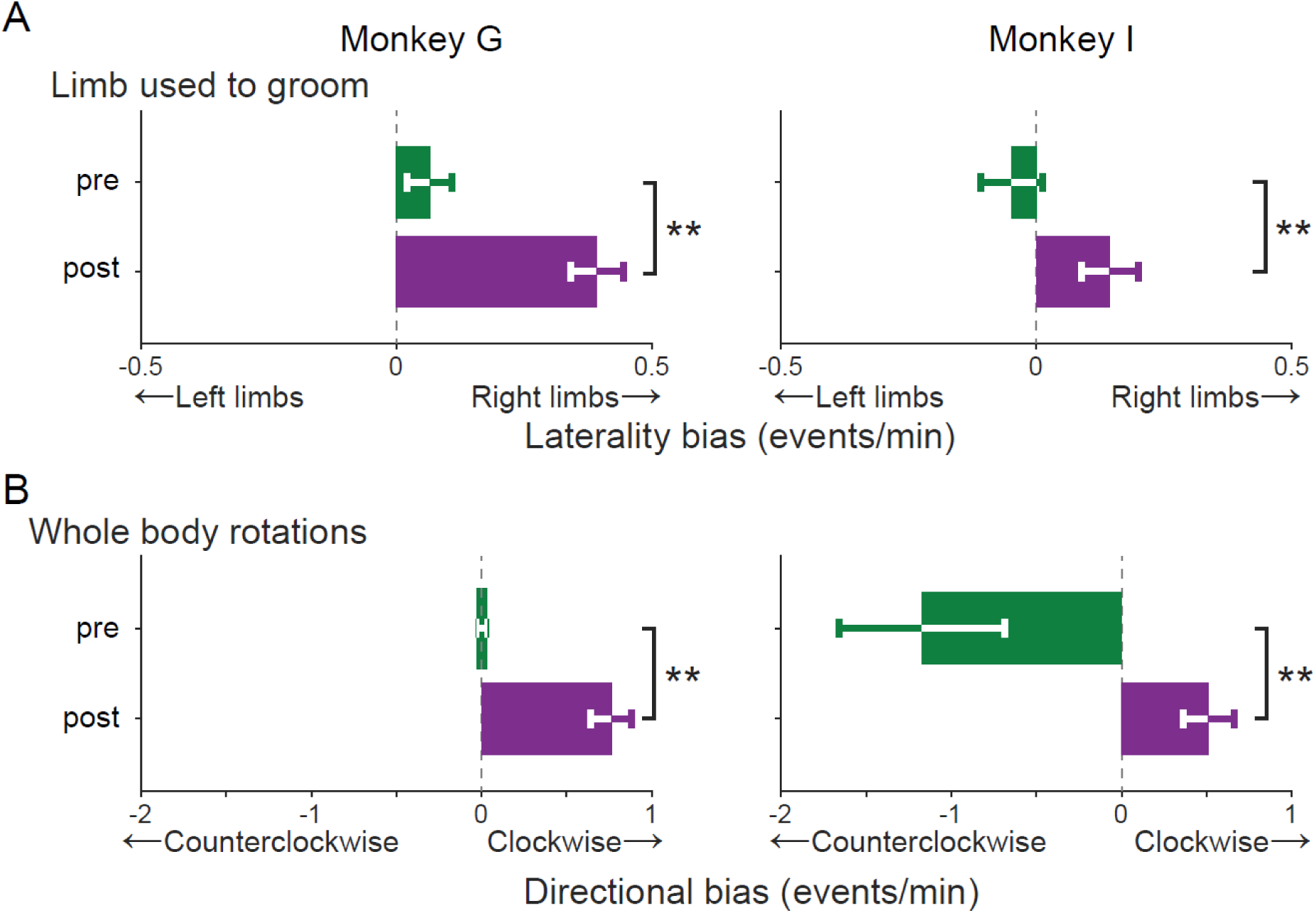
MPTP administration reduced use of the contralateral limbs and whole-body rotations toward the more affected side of the body. *A*. The rate of left limb use relative to right limb use during grooming behaviors before (green) and after (magenta) MPTP administration, shown separately for monkey G and I (left and right panels, respectively). *B*. The rate of clockwise whole-body rotations relative to that of counterclockwise rotations. Statistical comparisons were performed using the Wilcoxon rank-sum test (**p < 0.01).

**Figure S4.**
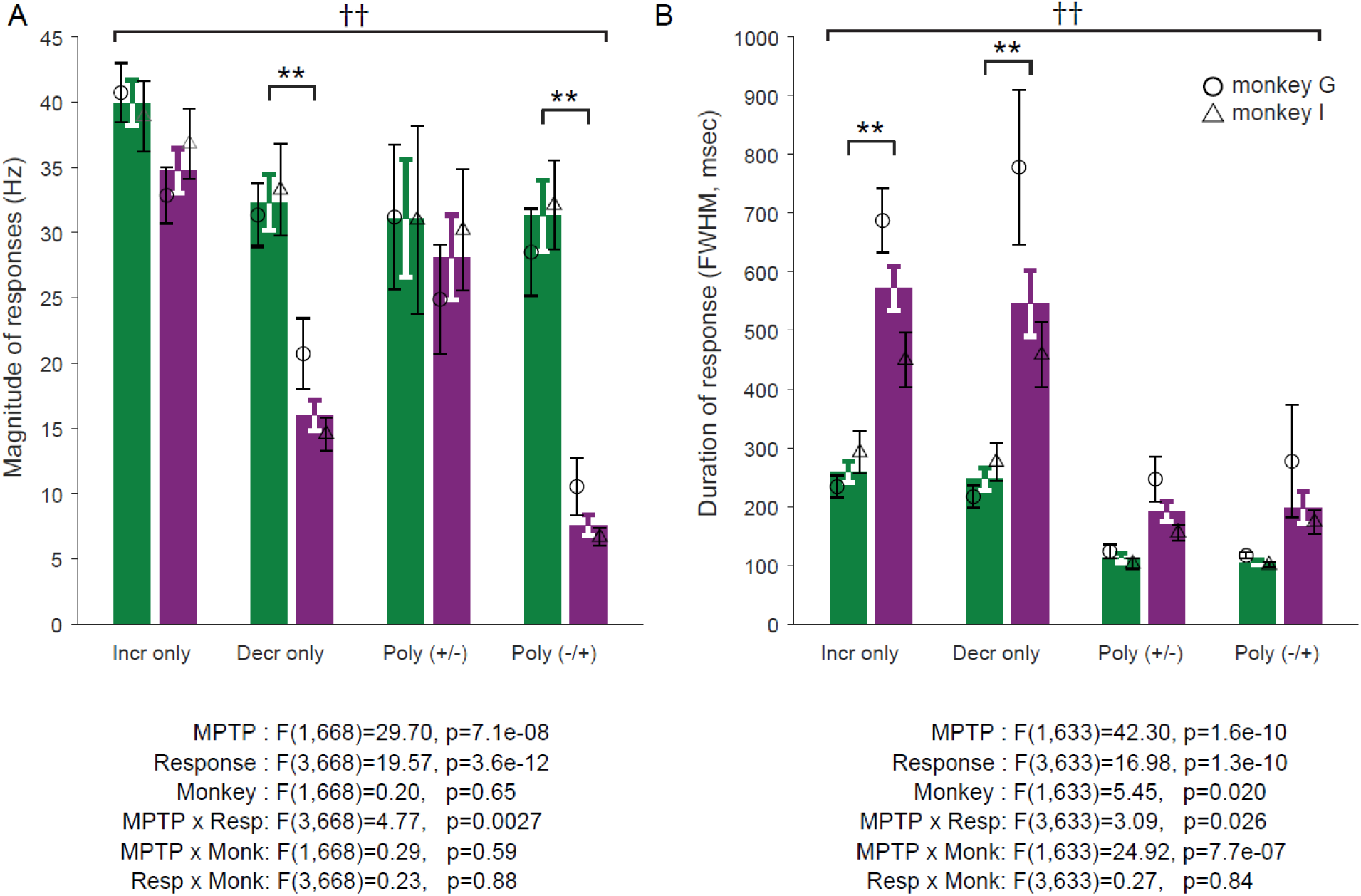
Effects of MPTP on response magnitude and duration as quantified from go-cue aligned SDFs. Results from this analysis were closely similar to those in measures taken from movement-aligned SDFs (Fig. 4D-E; following the same conventions). Results from 3-way ANOVA (MPTP × response type × animal) are shown below. *A*: The reduction in response magnitude was more severe in decrease-type responses (** p<0.01, Tukey’s test). *B*: Response durations were prolonged following MPTP administration, and that prolongation was more prominent for monophasic responses (** p<0.01, Tukey’s test).

**Figure S5.**
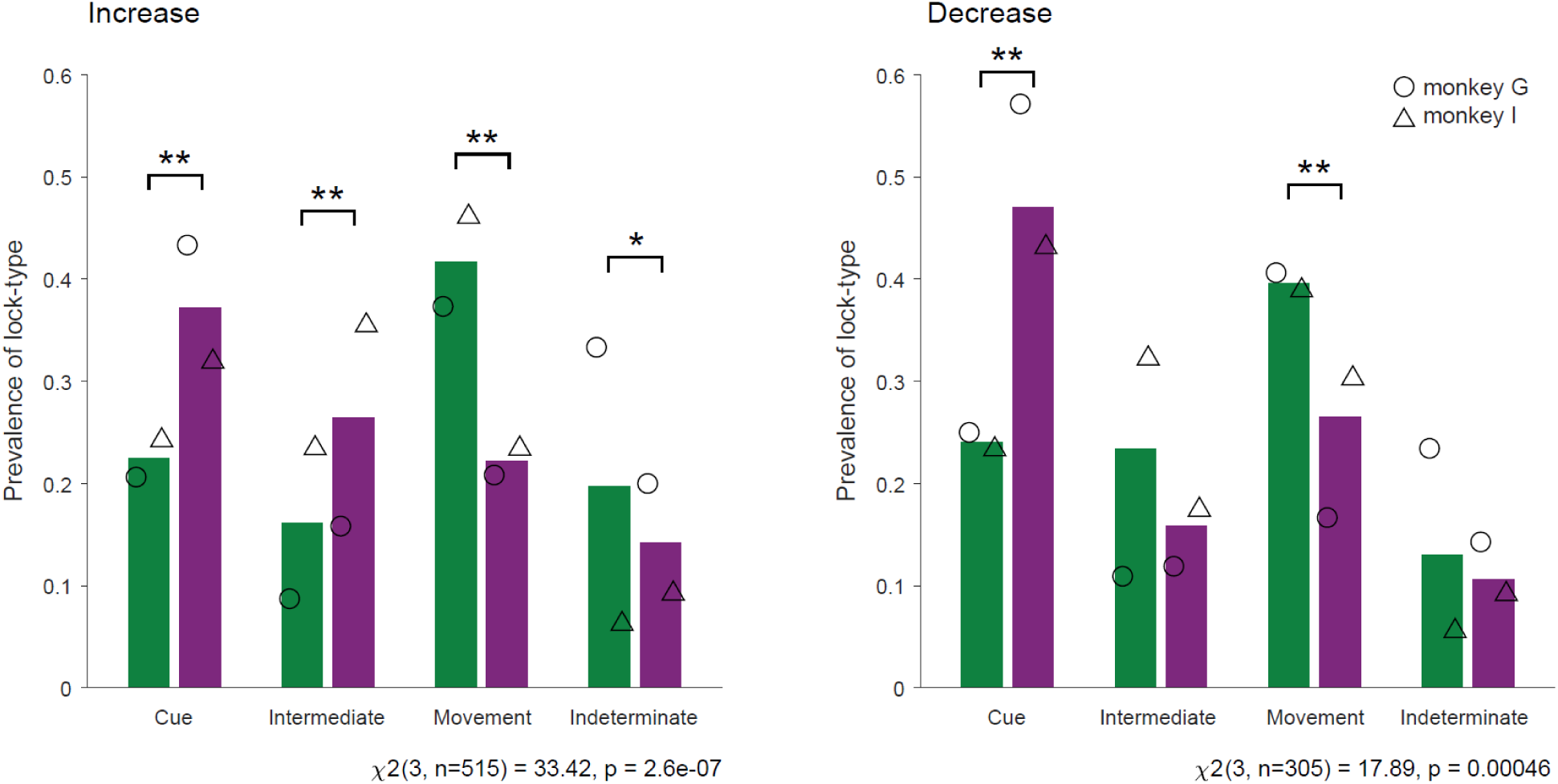
The increased prevalence of cue-locked responses following MPTP (i.e., Fig. 5D) was found in both increase-(*left*) and decrease-type response populations (*right*). Results from chi-square analysis are shown below figure panels (* p<0.05, ** p<0.01, adjusted residual analysis). *Open circles*: monkey G. *Open triangles*: monkey I

**Figure S6.**
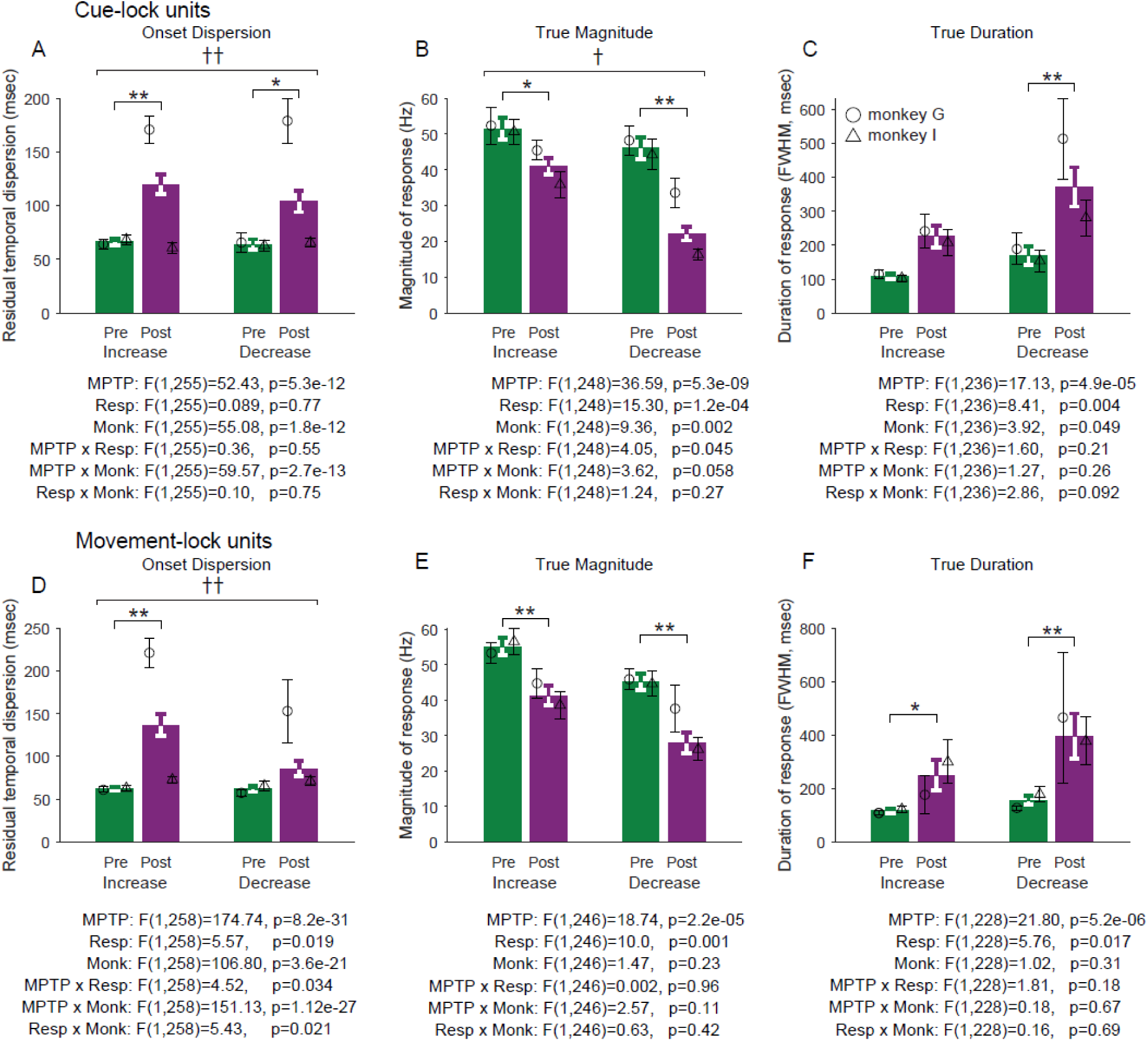
Effects of MPTP on jitter-corrected response metrics (dispersion, magnitude and duration; i.e., Fig. 6A-C) were similar in cue-locked (panels *A-C*) and movement-locked (panels *D-F*) response subtypes. Results from 3-way ANOVA (MPTP × response type × animal) are shown below.

**Figure S7.**
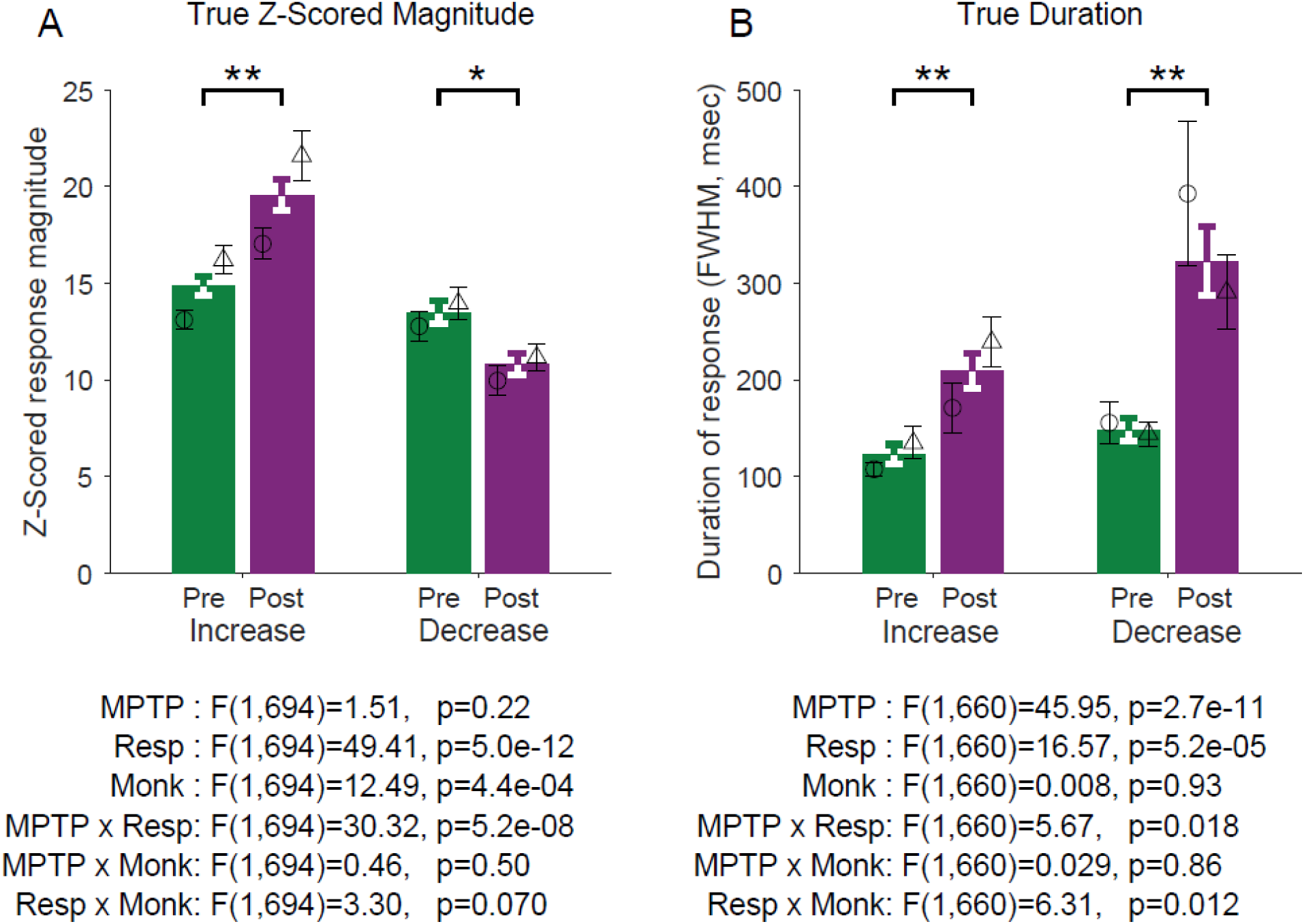
Effects of MPTP on Z-scored, jitter-corrected response metrics. *A*: The magnitude of increase-type responses was larger post-MPTP whereas the magnitude of decrease-type responses was diminished (* p<0.05, ** p<0.01, Tukey’s test). *B*: Response durations were prolonged following MPTP administration (** p<0.01, Tukey’s test). Results from 3-way ANOVA (MPTP × response type × animal) are shown below.

